# Prioritization of driver genes in cancer-associated copy number alterations identifies *B4GALT5* as a glycooncogene

**DOI:** 10.1101/2022.03.30.486261

**Authors:** Francesco Russo, Pranoy Sahu, Ilenia Agliarulo, Riccardo Rizzo, Matteo Lo Monte, Nicola Normanno, Silvia Soddu, Francesca Carlomagno, Alberto Luini, Seetharaman Parashuraman

## Abstract

Cancer is a disease resulting from aberrant communication between cells of a multicellular organism. The glycan coat that surrounds the cells is an important player in cellular communication. While altered cell surface glycans are known biomarkers for cancer, glycan biosynthesis itself has not been considered a potential oncogenic pathway. So, to understand the oncogenic potential of the glycan biosynthetic pathways we have analyzed the copy number alterations (CNA) of genes encoding for glycosylation regulators (glycogenes) in cancer genome datasets and identify novel glyco-oncogenes and glyco-tumor suppressor genes (TSGs). CNA of oncogenes and TSGs is an important cancer-associated genetic alteration that associates with worst prognosis. Nevertheless, identity of the driver genes in the copy number altered segments of the genome remains obscure in most cases. We developed a prioritization pipeline based on bioinformatic and experimental criteria to identify putative driver genes. In addition to correctly identifying several well-established oncogenes/TSGs, this pipeline discerns several novel oncogenes and TSGs, some of which are glycogenes. Further, among glyco-oncogenes there is an enrichment for glycosphingolipid biosynthetic pathway and *trans*-Golgi associated lysosomal sorting machinery and among glyco-TSGs there is an enrichment for early N-glycan biosynthetic enzymes. As a proof-of-principle we show that one of the identified glycooncogene *B4GALT5*, encoding a key enzyme in the glycosphingolipid pathway exhibits oncogenic property of promoting increased growth of hepatocellular carcinoma cells. Thus, this study identifies glycosylation pathways with oncoregulatory properties and opens up a new group of enzymes as potential therapeutic targets for cancer.

## Introduction

Cancer is a disease of the multi-cellular organisms resulting from the rogue behavior of one or more cells. These cells respond aberrantly to signals that coordinate cell growth and division leading to their uncontrolled growth and migration that ultimately affects vital functions [1], [2]. Thus, cancer can be viewed as a disease of aberrant cell communication. Glycan coat or glycocalyx forms the first layer of communication between cells. Many cell-cell and cell-matrix interactions are regulated by glycosylation of the cell surface receptors and/or ligands [3]. Underscoring the importance of glycans in cell-cell communication and in particular cancer, carcinogenesis is usually accompanied by changes in cell surface glycome [4], [5]. Indeed, glycans have been recognized as biomarkers of cancer from as early as 1960s [6], [7] and several cancer diagnostic kits based on the glycan profiling are currently used in clinic [8]. For instance, CA19-9 and CA72-4 are altered o-glycans that are used as biomarkers for several cancers and their presence is correlated POOR PROGNOSIS [8]. Further, studies have shown that the glycan changes accompanying carcinogenesis contributes to the development of cancer. For instance, the enhanced sialylation present on cancer cells has been shown to prevent key interactions involved in immune system mediated killing of cancer cells [9], [10], [11]. Further, the altered o-glycans on cancer cells have been directly linked to oncogenic features of cell growth and invasion [12], [13]. Along similar lines, mTORC2 has been shown to promote glycosphingolipid biosynthesis in liver cells required for tumor development [14]. In spite of these observations, a conceptual and mechanistic framework of how the glycans are altered in cancer cells and how they contribute to carcinogenesis is lacking.

Glycans are built in the secretory pathway, especially in the Golgi apparatus where majority of glycosyltransferases are localized. The human genome encodes for about 200 glycosylation associated enzymes whose expression levels and distribution in the determines the kind of glycans and hence the glycan coat of the cell [15]. By determining the glycan coat, the Golgi apparatus regulates cellular communication. A well-known example of this is the contribution of Golgi glycosylation to Notch signaling pathway frequently involved in cancer [16]. Identifying novel oncogenes or tumor suppressors whose products function at the Golgi to determine glycosylation may help in understanding the contribution of glycosylation to cancer.

A decade ago, an oncogene encoding for a Golgi localized protein, GOLPH3 was identified [17]. GOLPH3 is a protein adaptor that regulates sorting of glycosylation enzymes into COPI vesicles in the Golgi [18]. Indeed, altering GOLPH3 levels alters cell surface glycans, especially glycosphingolipids [19]. Studies had also shown that GOLPH3 plays an important role in secretion by promoting the exit of cargoes from the trans-Golgi network (TGN) [20]. Indeed, recent studies from our group [19] and others [21] have shown that the oncogenic role of GOLPH3 is linked to its function in regulating the localization of glycosylation enzymes in the Golgi, thus emerging as a prototypic oncogene regulating glycosylation. Following GOLPH3, genes encoding other Golgi localized proteins have been identified as oncogenes [22], [23], [24]. Whether they also act by regulating glycosylation is not clear. Nevertheless, these genes provide an important handle to understand the connection between Golgi processes, glycosylation and cancer. In order, explore systematically this connection we decided to look for potential novel oncogenes or tumor suppressor genes (TSGs) among genes involved in the regulation of glycosylation in the secretory pathway.

The changes in the activity of oncoproteins can be caused by several independent means. The most frequent mode of oncogenic activation (or inactivation in case of TSG) is point mutation that results in changes in the levels or activity of the protein. Another way to alter activity are gene fusions that by remove the regulatory controls acting on an oncoprotein to make it constitutively active. Levels of a protein (hence its total activity in the cell) can also be changed by copy number alterations (CNAs) and/or epigenetic alterations which are frequently observed in cancers [25]. The analysis of CNA or epigenetic alterations have lagged behind compared to analysis of cancer associated mutations mainly due to technological and/or informatic constraints, which have been recently overcome with the advent of next-generation sequencing and powerful algorithms to analyze the ensuing data [26]. The mutational landscape of the cancer, which has been extensively explored, is now considered to be saturated, at least in the case of common cancers [27] and such studies have not identified any oncogenic driver mutations in the glycosylation machinery of the cell [28]. Of note, germline mutations in proteoglycan processing enzymes EXT1 and EXT2 have been associated with multiple exostoses, but there are no reported somatic mutations in these genes that have been shown to act as oncogenic or tumor suppressor driver genes. On the contrary, the landscape of copy number alterations and epigenetic alterations associated with cancer is gaining ground and continues to yield novel candidate driver oncogenes and tumor suppressors. In fact, recently identified oncogenes encoding Golgi-localized proteins have been associated with the copy number alterations [17], [23]. In general, cancer-associated CNAs have been shown to have a better prognostic value in contrast to cancer-associated gene mutations [29]. Therefore, we decided to explore cancer genome-datasets to search for new oncogenes and/or TSGs from among the genes coding for the glycosylation machinery. Our study identified several potential oncogenes/TSGs from among the glycosylation associated genes. Further we found that pathways related to glycosphingolipid (GSL) biosynthesis and lysosomal sorting machinery at the TGN were enriched with new potential oncogenes and while N-glycan biosynthetic pathways was enriched with potential TSGs.

## Results

### Building a Golgi glycogene dataset

Human genome encodes about 25000-30000 protein-coding genes. An estimate of genes that contribute to glycosylation has ranged from 1-2% of the proteome (>800 genes) [30], includes enzymes involved in sugar-nucleotide biosynthesis, sugar transporters and glycosylation enzymes present in the Golgi and others. From a literature search and manual curation, we obtained a dataset of 479 proteins that likely controls glycosylation reactions in the Golgi, hereinafter referred to as glycogenes. Glycogenes included glycosylation enzymes and glycosidases (236; including Glycosylphosphatidylinositol (GPI) anchor biosynthesis and endoplasmic reticulum (ER)-localized dolichol sugar biosynthases), channels and transporters (40), membrane trafficking machinery (169), components of lipid trafficking and metabolism (34; including phosphoinositide metabolism) (Fig. 1A; Table S1). These genes were distributed all across the genome with no particular concentration in any chromosome or cytogenetic band (Fig. S1A,B). Next, we investigated whether some glycogenes may have oncogenic features. So, we analyzed the published focal copy number alterations (FCNAs) and found that these FCNAs encompassed regions where several glycogenes were located (Fig. S1C). We also noted that expression of several glycogenes were altered in cancers and their expression levels correlated with the survival of patients (Fig. S1D). Subsequently, we set-up a pipeline to identify potential glyco-oncogenes and glyco-TSGs among glycogenes displaying CNAs.

**Figure 1:**
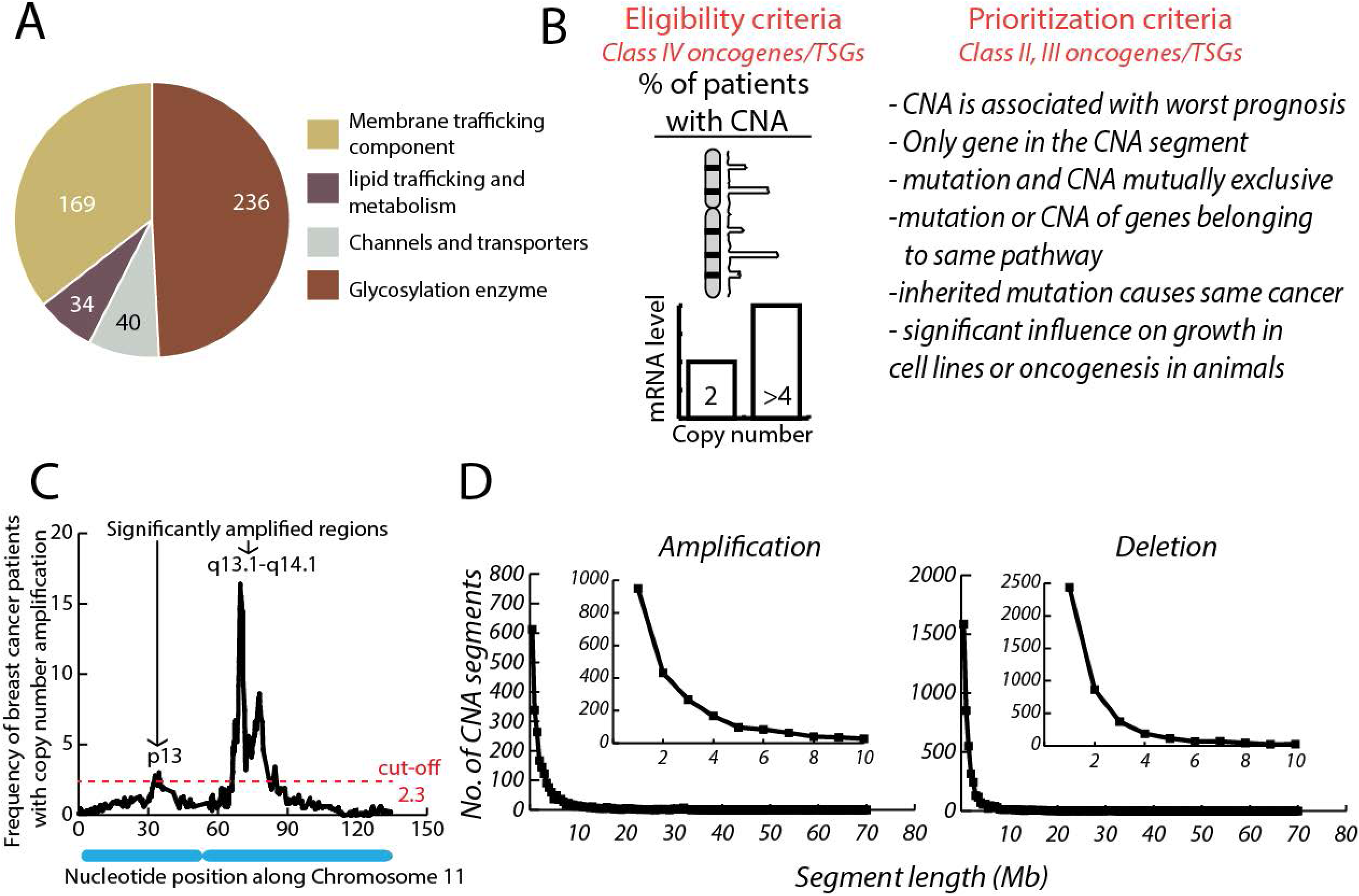
Development of pipeline to identify oncogenes/TSGs based on CNA. **A**. Classification of glycogenes analyzed in the study. **B**. Pipeline to identify and prioritize genes with significant CNA in the cancer genomes. The eligibility criteria first identified genes with significant CNA in cancer genome and then those with a corresponding statistically significant alteration in the expression level are classified as Class IV oncogenes/TSGs. Based on the listed criteria, the Class IV genes were then prioritized and classified as Class II or III genes. **C**. A chromosomal level threshold for frequency of CNA was identified (see Methods) and the chromosomal segments above the threshold we considered to have statistically significant CNA. An example of thresholding on Chromosome 11 in Breast invasive carcinoma is shown along with the two segments (p13 and q13.1-q14.1) that showed statistically significant amplification among the patients. **D**. the length distribution of identified CAN segments is shown with most of them less than 5Mb in length.

### Pipeline to prioritize candidate glyco-oncogenes/glyco-TSGs

To identify and prioritize candidate glyco-oncogenes we focused on CNAs for the above-described reasons (see introduction). For a gene to be considered an oncogene based on CNA, a set of criteria have been proposed [31]. The minimal criteria were that the gene should exhibit a statistically significant CNA in cancer genomes and a corresponding alteration in its expression level. Genes that pass these criteria were further prioritized based on other characteristics that can potentially contribute to oncogenesis. These included clinical (positive correlation between CNA of a gene and poor prognosis), biological (experimental evidence of its contribution to oncogenesis) and bioinformatic (mutual exclusivity of mutation and CNA of a gene, being the only gene in the copy-number altered segment of the genome and others) parameters (see below for details). Genes that exhibit only the minimal criteria were termed as Class IV oncogenes, while depending on the number of other criteria that they satisfy they were classified as Class III or II oncogenes. In addition to all these, if measures to counter the gene activity (or its loss in case of TSG) resulted in favorable clinical outcomes in patients with CNA of the gene, it was classified as a Class I oncogene. Of the 70 described CNA based oncogenes only 2 fulfilled all the criteria to be considered a Class I oncogene [31]. Our pipeline here mirrored these criteria (Fig.1B) where we first start with the identification of chromosomal segments that are significantly altered in cancer genomes, followed by selecting genes encoded in these segments whose expression levels match their CNA. This is the minimum criteria to make a gene eligible for consideration. These genes were further tested for the other mentioned criteria to arrive at a stratified priority list of potential oncogenes/TSGs.

### Identification of chromosomal segments with significant alteration in copy number in cancers

To identify chromosomal segments with significant alteration in their copy number in cancer, we analyzed the publicly available The Cancer Genome Atlas (TCGA) datasets provided by cBioportal (www.cbioportal.org) [32]. The datasets consisted of genomic, transcriptomic, and clinical data of 32 different cancer types (Table 1). The number of patients ranged from a minimum of 36 patients in case of Cholangiocarcinoma to a maximum of 1080 patients in the case of Breast Invasive Carcinoma for a total number of 10845 patients. For each cancer type, the copy number data for 24776 genes were obtained as GISTIC (Genomic Identification of Significant Targets in Cancer) scores (calculated by the GISTIC2.0 algorithm [33]. These datasets have previously been characterized and shown to be broadly representative of the general (U.S) population. [29].

To identify the copy number altered segments of a chromosome, we have focused only on those genes that showed true amplification (copy number >4) or homo deletion (both copies deleted). The frequency of CNA of genes in each of the 32 cancer types were plotted along the chromosomal length. A background threshold was calculated based on the distribution of these observed frequencies, to identify genes with significantly frequent CNA in a chromosome (see Methods). Then the chromosomal segments representing CNAs were identified based on the following criteria (Fig.1C): the chromosomal segment is composed of a contiguous set of genes that show an above threshold frequency of alteration in the same direction (amplification or deletion) and are altered in the same set of patients. Finally, only those copy number altered segments with the length of 1kb to 100Mb [34] was considered for further analysis so as to focus on FCNAs. Based on these criteria, we identified 6920 copy number altered segments - 2413 amplified and 4507 homo deleted - across the cancer types. Uterine cervical Carcinoma (UCEC) had the maximum number of amplified segments (186) and uveal melanoma the minimum number of amplified segments (10). In case of deleted chromosomal regions, prostate adenocarcinoma had the maximum (403) and kidney chromophobe the minimum number of segments (37) (Fig.S2A). In most cancer types the number of statistically significant amplified and deleted chromosomal segments identified had near linear correlation except for prostate adenocarcinoma where we found more deleted segments (403) compared to amplified segments (53) (Fig. S2B). A similar linear correlation between the distributions of amplified and deleted chromosomal segments across the chromosomes was also observed (Fig.S2C). Most of the copy number altered segments were less than 5Mb in size (Fig.1D) and median length of the CNA segments in cancers ranged from about 0.27 Mb in case of diffuse large B-Cell lymphoma (DLBC) to 2.8 Mb in case of lower grade glioma (LGG) (Fig.S3A). Further, the frequency of a segment was inversely proportional to the length of the CNA segment (Fig.1D) similar to what was described previously [34]. We then checked if the identified regions showed significant overlap with the already published copy number altered regions in these cancers. In 18 cancer types where published data were available, we identified a median of 76% of the published amplified segments and 64% of the published deleted segments in our study (Fig.S3B). The only outlier was pancreatic adeno carcinoma (PAAD) where only 15% of the published deleted segments was also identified by us (Fig.S3B; Table S2). We also checked the overlap of our data with another published analysis of TCGA datasets for CNA [35] and found our analysis identified about 90% of the published significantly amplified genes (Fisher test: and 74% of the published significantly deleted genes from this study Thus, this significant overlap with the published results suggests that our method was robust enough to capture the copy number altered segments associated with cancers.

We next analyzed the pan-cancer amplification/deletion of the chromosomal segments (defined here as chromosomal segments with statistically significant CNA in at least 10 cancer types). We identified a total of 103 and 102 cytobands that were amplified or deleted respectively in more than 10 cancer types (Fig.S4 and Table S3). These segments included the well-established oncogenes EGFR and Myc and TSGs RB1, PTEN and TP53. As expected, the pan-cancer amplification and deletion segments tended to be mutually exclusive. Further, the CNA segments were also often found in the end of the chromosomes. While some chromosomes like Chromosome 14 and 21 barely had any pan-cancer CNA segments, others like Chromosome 1 had several (Table S3). There was no correlation between the number of pan-cancer amplified and deleted segments in a chromosome (Fig.S5A) while there was a weak linear correlation between the total number of pan-cancer CNA segments identified and the length of a chromosome (Fig.S5B). Next, we compared them with the published list of pan-cancer FCNAs [34]. Of the published set of FCNAs nearly 40% were identified also in our pan-cancer list (present in more than 10 cancer types). When threshold of 10 was relaxed to include at least 2 cancer types, nearly all the published FCNAs were retrieved (data not shown).

The median frequency of patients showing copy number amplifications was 3.5% with minimum being 0.25% in the case of 15q21.3 in THCA and maximum being 42.3% in the case of 3q26.32-q29 in case of lung squamous cell carcinoma (LUSC). The median frequency for copy number deletions was 2.2%, with the minimum being 0.3% in the case of several chromosomal segments in glioblastoma (GBM) and the maximum being 31.03% in the case of chromosomal segment 9p21.3 in mesothelioma (MESO). Most of the CNA were in less than 5% of the patients (71.5% for amplification and 85.2% for deletions) with a peak at 3% of the patients for amplification and 2% in case of deletion (Fig.S5C). Nearly 99% of the identified CNA were present in less than 15% of the patients in each cancer type (Fig.S5C). These numbers in general reflect earlier published frequencies of copy number alterations [34].

Thus, we were able to identify copy numbered altered segments in the cancer genome with characteristics-length, frequency of amplification and overlap with the previously published results that match with that expected of FCNAs.

### Identification of potential oncogenes based on CNA

The eligibility criteria for a gene to be classified as an oncogene/TSG based on CNA are 1) it shows a statistically significant CNA among cancer patients and 2) there is a corresponding alteration in the gene expression [31]. The genes that possess these two characteristics are classified as Class IV oncogenes (Fig.1A). So we analyzed genes with significant CNA (see above), to identify those with statistically significant change in expression levels (p<0.05 in a Wilcoxon-Mann-Whitney test with Benjamini Hochberg correction) and in the appropriate direction i.e., an increase in case of gene amplifications and a decrease in case of gene deletions. The cut-off for expression level changes was set to a fold change of >1.2 for oncogenes and <0.8 for TSGs. This resulted in the 7737 Class IV oncogenes and 4865 Class IV TSGs (see above) being selected across all the cancer types (Fig.2A). For amplification, the number of genes per cancer type varied from a minimum of 4 genes (in kidney chromophobe) to a maximum of 2705 genes (in ovarian cancer). In case of deletion, the minimum was 1 gene (in kidney chromophobe) and the maximum was 1513 genes (in prostate adenocarcinoma).

**Figure 2:**
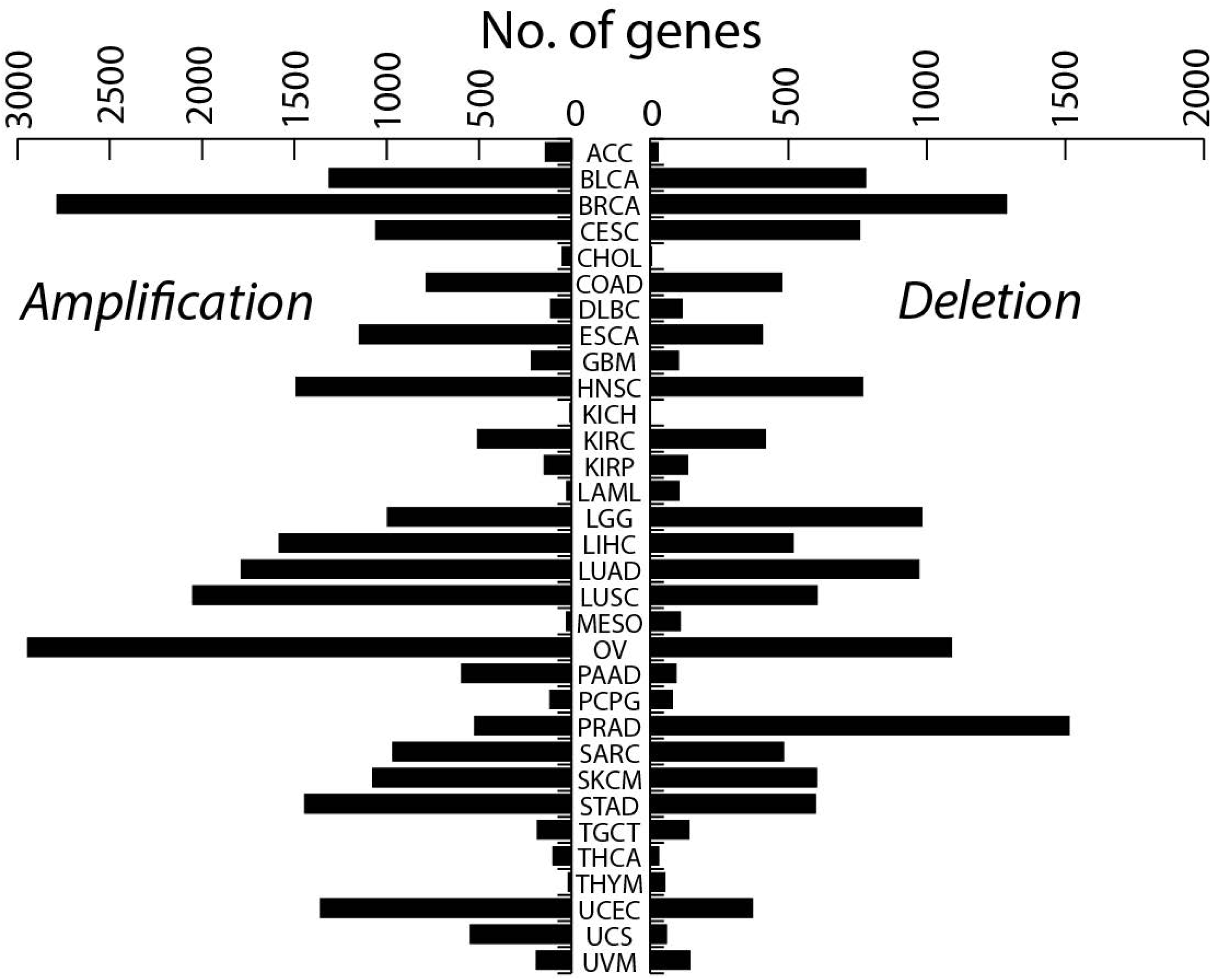
Genes with significant CNA. The number Class IV oncogenes and TSGs identified in each cancer type is plotted.

Among the previously identified 88 known amplification-based oncogenes (See Table S4), 74 were identified as Class IV genes in our analysis while of the 41 known tumor suppressors (based on CNA; See Table S4) 27 were identified as Class IV genes. The low overlap in case of TSGs might be related to the fact that we included only homodeleted genes in our analysis. Further, several mutation-based oncogenes and tumor suppressors were present in the list of Class IV oncogenes/TSGs. We also noted that not all the genes present in a chromosomal segment with CNA had significantly altered expression. This suggests that mechanisms other than CNA, such as methylation of promoters and/or mRNA stability, can affect the final level of expression of a gene.

### Prioritization of potential oncogenes/TSGs

Next to prioritize the Class IV oncogenes/TSGs we analyzed the following characteristics of these genes:

*Clinical:* 1. correlation of CNA with patient survival,

*Bioinformatic:* 2. presence of more than one Class IV gene in a chromosomal segment with CNA, 3. mutual exclusion of mutation and CNA in patients, 4. documented evidence for inherited mutation in the gene leading to cancer, 5. more than one gene from the same pathway is amplified or mutated in the same cancer type,

*Biological:* 6. overexpression (oncogenes) or downregulation (TSG) of the gene of interest promotes cell growth in vitro, 7. downregulation of a Class IV oncogene in a cell line where it is amplified reduces cell growth/viability and 8. finally transposon mediated deletion/inactivation of a Class IV TSG promotes carcinogenesis in animals.

Each of these characteristics was assigned a point and those Class IV genes that acquire 1 or 2 points were classified as Class III oncogenes/TSGs and those that get 3 or more points were classified as Class II oncogenes/TSGs [31].

To evaluate whether the patients with CNA of a gene had a worse survival compared to those without, we used the Kaplan-Meier estimator. Only those genes that show CNA in at least 5 patients were considered for analysis and tests were done in cancer-stage matched manner (when data was available; see Methods). Those genes where the patients with CNA showed a significantly worse survival compared to the rest were assigned a point. Among the amplified and overexpressed genes, we identified 2652 genes whose CNA showed significant correlation with worst prognosis. As for other criteria, 312 genes were found as a unique gene in the amplified chromosomal segment, 133 showed mutual exclusion between mutation and CNA, 353 genes had at least one mutated or amplified gene in the same cancer type that belonged to the same Reactome pathway, 169 genes when knocked down by siRNA reduced cell growth/viability in cell lines where the gene was amplified, 40 genes were found to promote cell growth in a published overexpression screen and finally inherited mutations in 5 genes have been described to cause similar type of cancer where they were found amplified.

Among the deleted genes, 826 genes showed significant correlation with worst prognosis (*Clinical relevance* is 1). As for other criteria, 797 genes were found as a unique gene in the deleted segment, 189 showed mutual exclusion between mutation and deletion, 955 genes had at least one mutated or deleted genes belonging to the same Reactome pathway, 455 genes when knocked down by siRNA promoted cell growth/viability in cell lines derived from the same cancer type where they were found deleted, 191 genes were found to promote oncogenesis following insertional mutagenesis screens in mice and finally inherited mutations in 12 genes have been described to cause similar type of cancer where they were found deleted.

Altogether, based on the points scored we were able to classify 2510 genes as Class III oncogenes and 39 as Class II oncogenes. As for TSGs, 1635 were found to be Class III TSGs and 147 were found to be Class II TSGs (Fig.3A, Table S5).

### Validation of the prioritized list of oncogenes and TSGs

The prioritization procedure described above had led to a list of potential oncogenes and TSGs whose CNA contributes to oncogenic development. In order to understand if this prioritized list is indeed enriched in oncogenic regulators, we carried out a series of investigations. First, we checked the overlap of our priority list with cancer census genes list from COSMIC. Of the 723 genes present in this gene list, 140 were classified as oncogenes and 135 as TSGs in our priority list, which was a highly significant overlap (Fisher Exact Test; *p-value* < 10^−15^). Next, we checked how many of the known CNA based oncogenes and TSGs (as described in COSMIC database and literature; Table S4) were identified in our priority list. Of the 88 known amplification-based oncogenes, 50 were classified as Class II or III oncogenes in our priority list (Fisher Exact Test; *p-value* < 10^−16^). Similarly of the 41 CNA based TSGs, 6 were found in our list (Fisher Exact Test; *p-value* < 0.5). Thus, nearly 50% of the known CNA-based oncogenes and 10% of known CNA-based TSGs were also identified in this analysis (Fig.3B-C). The list included major known oncogenes/TSGs like *EGFR, RICTOR, MET, CDK6* and *CCNE1* classified as Class II oncogenes and *APC, BRCA2, PTEN, CDKN2A* and *RB1* classified as Class II TSGs. Of note, several known mutation-based oncogenes/TSGs were also present in our priority list of CNA-based oncogenes/TSGs. Next, we investigated the GO processes that are enriched in this prioritized list of genes. We found that among the oncogenes, statistically significant enrichment was found for mitochondrial translation elongation and DNA replication. While among the TSGs, positive regulators of apoptosis, negative regulators of cell proliferation and regulators of cell cycle arrest were enriched. While processes enriched among TSGs were expected, those among oncogenes were slightly surprising. We expected cell cycle regulators and growth signalling pathways but surprisingly the top hit was mitochondrial translation elongation. Given the important role of mitochondrial biology in carcinogenesis and application of mitochondrial translation inhibitors to target cancers, especially cancer stem cells [36] the finding here acquires importance. Testing CNA of relevant genes in mitochondrial translation may help direct antibiotic based therapeutic measures to relevant patients. The enrichment of DNA replication machinery suggests the contribution of CNA based oncogenes to increased cell division. These analysis in sum suggests that the prioritization pipeline developed here seems robust and dependable.

While prioritized list of genes obtained by could identify several known oncogenes/TSGs and is enriched in regulators of known oncogenic mechanisms, we wanted to know if this list also points to new unknown regulators that might lead to identification of novel oncogenic regulators/pathways. This we thought would provide support to the novel glycogenes based oncogenic regulators that we wished identify. In order to do this, we based our analysis on the finding that the interactors of the known oncogenes are enriched in novel oncogenic regulators [37]. So, we analyzed the enrichment of direct interactors (https://string-db.org/) of COSMIC cancer census genes among the identified oncogenes and TSGs. We found 795 of these interactors in the list of prioritized oncogenes and 553 of them in the prioritized list of TSGs. The overlap of the datasets of cancer census gene interactors and the list of oncogenes/TSGs showed a strong statistical significance (Fisher Exact Test; *p-value* < 10^−16^). Thus, we conclude that the prioritized CNA based oncogene/TSG list that we have obtained is enriched in the several novel oncogenes/TSGs.

### Glyco-oncogenes and glyco-TSGs

Among the Class II and Class III oncogenes/TSGs identified through our pipeline, we found 80 Class III glyco-oncogenes and 52 Class III glyco-TSGs (Fig.3D). There were no Class II glyco-oncogenes or glyco-TSGs. Three genes were present in list of both glyco-oncogenes and glyco-TSGs – *SEC22B, B3GALNT2 and ACBD3*. As a fraction of all Class II/III oncogenes/TSGs identified, glyco-oncogenes were enriched (> 5% of the identified genes) in ACC, COAD and PAAD, while glyco-TSGs were enriched in THYM, UCS, PRAD and UCEC (Fig.3E). Of note, the glycogenes constitute less than 2% of the total of number genes (24776) analyzed.

**Figure 3:**
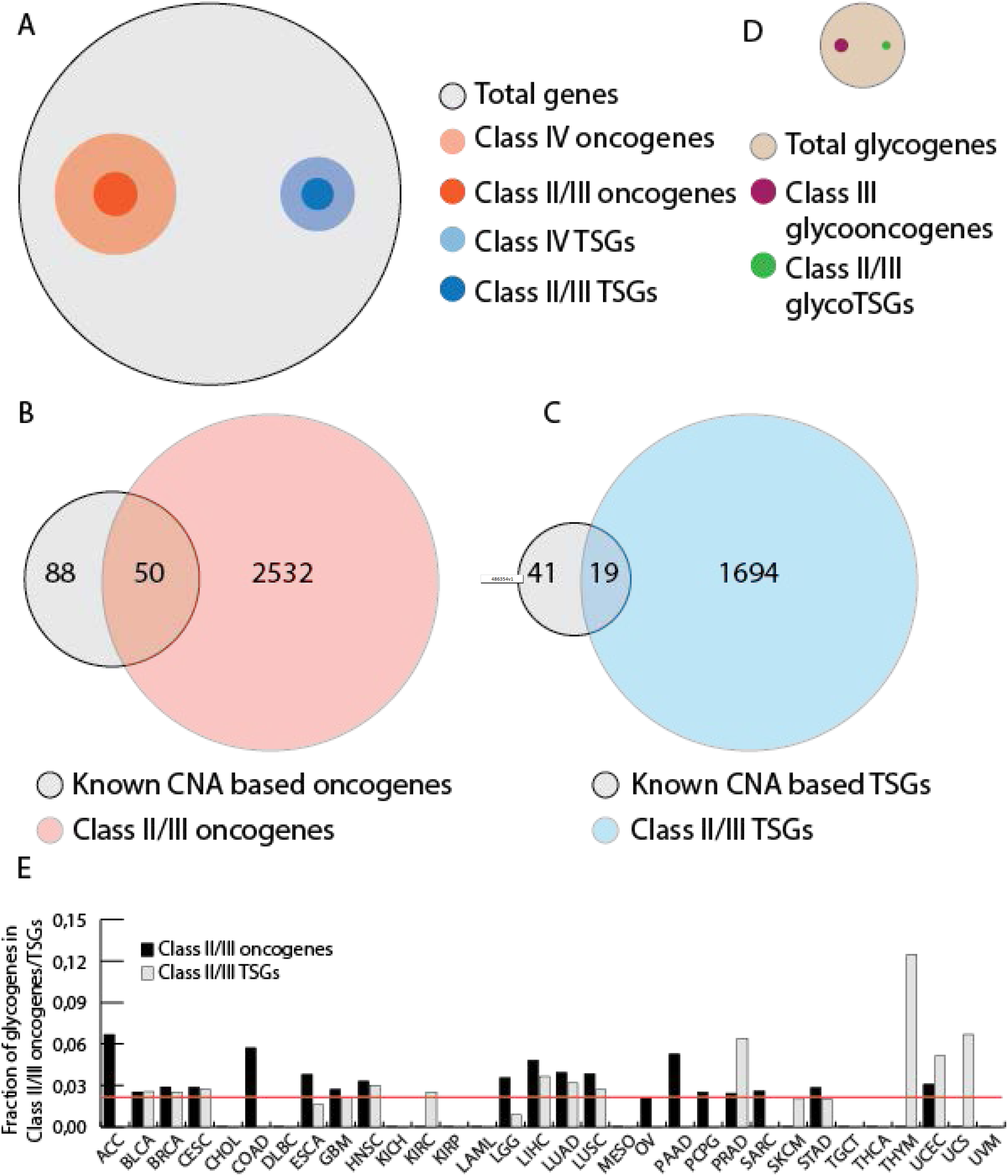
Prioritization of Class IV oncogenes and TSGs. **A**. Venn diagrams showing the relationship between total number of genes analyzed, the number of identified as Class IV oncogenes or TSGs and the number of Class II/III oncogenes or TSGs identified. **B-C**. Venn diagrams of total number of Class II/III oncogenes and TSGs identified with respect to the known CNA based oncogenes and TSGs. **D**. The same as A but refers only to the glycogenes. **E**. The number of Class II/III glyco-oncogenes and glyco-TSGs across cancer types is plotted as the fraction of identified Class II/III oncogenes and TSGs. The red line indicates the fraction glycogenes in the total number of genes analyzed in this study.

To understand further the glyco-oncogenes and glyco-TSGs, we investigated if there are any differences in the pathways to which they belong. So, we analyzed for the enrichment of KEGG (Kyoto Encyclopedia of Genes and Genomes) pathway among these genes. Glyco-oncogenes were enriched in lysosome, GSL biosynthesis (lacto/neolacto and ganglio series), glycosaminoglycan biosynthesis (chondroitin sulphate /dermatan sulphate and heparin sulfate/heparin), GPI anchor biosynthesis and other types of O-glycan biosynthesis (Notch glycosylation) (Fig.4A). While glyco-TSGs were enriched in N-glycan biosynthesis, glycosaminoglycan biosynthesis (heparan sulphate/heparin) and GPI anchor biosynthesis (Fig.4A). The enrichment analyses showed a distinguishing feature of glyco-oncogenes and glyco-TSGs with GSL biosynthetic genes, lysosomal sorting determinants and Notch glycosylation enriched only among glyco-oncogenes and N-glycan biosynthetic genes enriched among TSGs. GAG biosynthetic genes and GPI anchor biosynthetic genes were enriched in both glyco-oncogenes and glyco-TSGs. When dissected further, it was seen that enrichment of GAG biosynthetic pathway in glyco-oncogenes included the early part of the GAG biosynthetic pathway (*B4GALT7* and *XYLT1*) that is common to all 3 types of GAG biosynthesis while among the enriched GAG genes in glyco-TSGs all the genes present belonged specifically to heparin sulphate biosynthesis. In the case of GPI anchor biosynthesis, the glyco-oncogenes were mainly involved in the biosynthesis of the core saccharide unit of the GPI anchor while glyco-TSGs were involved in the modification of the core saccharide (see below). Next, we explored further the link between these enriched pathways and oncogenesis.

**Figure 4:**
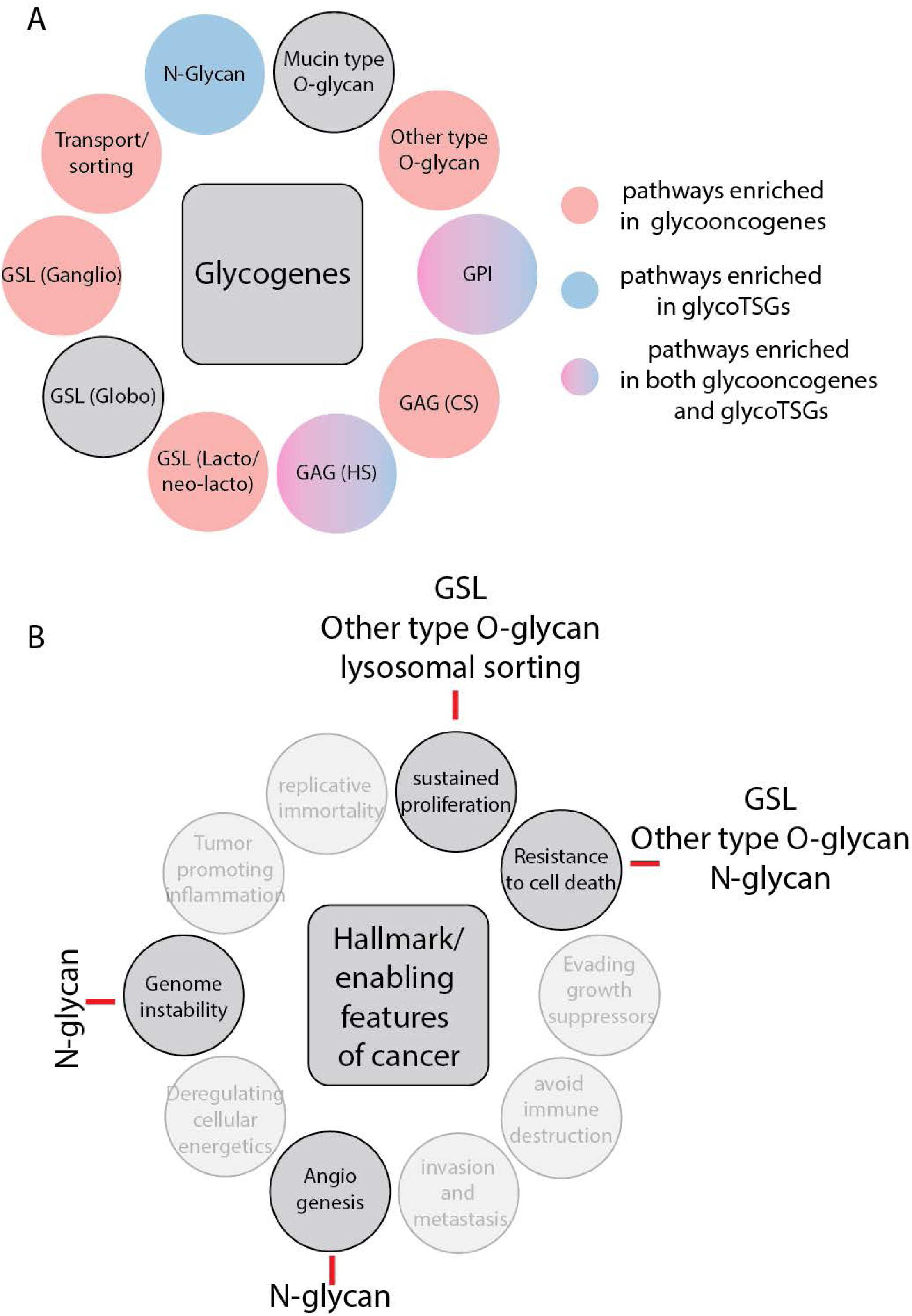
Characterization of Class II/III glyco-oncogenes and glyco-TSGs. **A**. the glycan (transport) pathways encoded by the glycogenes is shown along their enrichment in glycogenes and glyco-TSGs respectively. **B**. Hallmark and enabling features of cancers are shown. When an identified Class II/III glycooncogene or glycoTSG belonging to indicated glycan pathways show interaction (as identified through STRING database) to genes contributing the indicated features of cancer, this interaction is shown by a red line.

### Glycosphingolipid (GSL) biosynthetic pathway

The genes of the GSL pathway that were classified as potential oncogenes include *B3GNT2, B3GNT3, B3GNT5* and *FUT4* involved in the biosynthesis of lacto-neolacto series GSLs and *B4GALNT1, ST6GALNAC6* and *SLC33A1* involved in the biosynthesis of ganglio series GSLs. In addition, *B4GALT5* encoding for lactosylceramide synthase an enzyme that produces a precursor for both lacto/neolacto and ganglio series, *GOLPH3* encoding a recently identified controller of B4GALT5 [19] and *ORMDL3* encoding a regulator of sphingolipid biosynthesis [38] were also present in the list. The genes were not consistently found in any cancer type. To understand how these proteins may contribute to carcinogenesis, we studied their interaction with the cancer census genes (COSMIC database) [39]. We identified an interaction of B4GALT5 and B3GNT5 with mucins (MUC4, MUC16 and MUC1). The mucins have been linked to the promotion of growth factor signaling, inhibition of cell death pathways and invasion which are three hallmark properties of cancer [40]. Similarly, the GSL production has also been linked to the promotion of growth factor signaling and invasion [41]. In addition, SLC33A1 (Acetyl-CoA transporter, involved in acetylation of GSLs) has been identified as an interactor of ATM, a kinase involved in DNA repair signaling [42]. Whether this interaction has a functional significance for DNA repair signaling requires experimental validation. Thus, GSLs can potentially contribute to three hallmark properties of cancer – promotion of growth, inhibition of cell death and invasion (Fig.4B).

### Lysosomal sorting

The genes of the lysosomal sorting machinery that were classified as potential oncogenes include *GGA2, GGA3, AP1S1, AP4S1, AP4M1, AP3M2, CLTC, GNPTG* and *LAPTM4A*. Most of the genes identified include the sorting machinery acting at the TGN. GNPTG is N-Acetylglucosamine-1-Phosphate Transferase Subunit Gamma that is involved in the production of Mannose-6-phosphate signal for lysosomal sorting and LAPTM4A is a lysosomal localized protein which has also been found in the Golgi apparatus [43]. The exact role of the protein is not clear, but its paralog LAPTM4B has been shown to be overexpressed in some cancers [44]. Three of this pathway genes (*AP4S1, GGA2, GNPTG*) were found amplified in breast cancer. *CLTC* (Clathrin heavy chain subunit) has already been implicated in carcinogenesis as fusion partner for *ALK* and *TFE3* [45],[46]. In addition, it has also been shown to regulate kinetochore stability [47], to regulate p53 mediated transcription [48] and to promote stability and nuclear translocation of HIF1alpha required for expression of VEGF [49]. When we analyzed the interactors of these potential oncogenes, we found that they interact with 3 others present in cancer census gene list – *EPS15, MET* and *RABEP1*. Interaction of CLTC with EPS15 is required to regulate the endocytosis of several growth factor receptors [50] and also for transport from TGN [51] and the interaction of GGA3 with MET has been linked to promote recycling of MET and prevent its sorting towards degradation [52]. Similarly, GGA2 has been implicated in sorting of EGFR [53]. GGA2 and GGA3 also interact with RABEP1 or Rabaptin that is also involved in promoting sorting into recycling endosomes [54]. Thus, the sorting machinery embedded among this list of potential oncogenes seem to regulate endocytosis and sorting to recycling endosomes which controls growth factor signaling (Fig.4B). LAPTM4A has also been shown regulate GSL biosynthesis [43] and so it can potentially act on oncogenesis through regulation of GSL biosynthesis (see above).

### GAG

The genes of the GAG biosynthetic pathway that were classified as potential oncogenes include *B4GALT7, XYLT1, EXT1, CHST12* and *CHSY1*. B4GALT7 and XYLT1 are involved in the initiation of GAG biosynthesis producing the first two sugar units of the tetrasaccharide unit that is the precursor for the biosynthesis of all types of GAGs. While *CHST12* and *CHSY1* are involved in chondroitin/dermatan sulfate biosynthesis and *EXT1* in heparan sulfate biosynthesis. Other GAG biosynthetic genes, specifically those involved in heparan sulfate biosynthesis were also found to be enriched among glyco-TSGs. All of these genes *EXT2, EXTL3, HS3ST3A1* and *NDST3* were involved in the distal part of HS biosynthetic pathway. *EXT2* and *EXTL3* are involved in the extension of alternating disaccharides that form constitute the HS chains while *HS3ST3A1* and *NDST3* are involved in the sulfation of the heparan disaccharide chain. Interactors of the GAG genes from the cancer census genes include glypicans-3 and 5 and syndecan-4, all of which have been shown to act mainly as tumor suppressors [55], [56], [57] and in some cases also as oncogenes [58], [57]. So, it is not clear how really GAG genes may contribute to oncogenesis. The germline inactivating mutations in *EXT1* and *EXT2* caused multiple exostoses [59], suggesting that they may actually act as tumor suppressors but more experimental analysis is required to test the hypothesis.

### N-glycan

Among the N-glycan biosynthetic genes enriched among the glyco-TSGs there were *ALG1, ALG9, ALG11, ALG12, MAN1A2, MAN2A1, MGAT5, STT3A* and *STT3B. ALG1, ALG9, ALG11, ALG12, STT3A* and *STT3B* are involved in the biosynthesis and transfer of dolichol-linked oligosaccharides to the proteins in the ER. *MAN1A2, MAN2A1* and *MGAT5* are involved in the processing of N-glycans in the cis-medial Golgi. Among the interactors of these gene products, we found CUX5, GOLGA5 and RPN1. The latter two act as gene fusions with known oncogenes to promote oncogenesis [60], [61] so their direct contribution to oncogenesis has not been observed. *CUX1*, a homeobox gene on the other hand has been shown to act as a tumor suppressor as well as an oncogene. Its absence has been shown to resistance to apoptosis and increase in DNA instability [62]. Paradoxically in some tumors it has also been shown promote growth factor signaling and angiogenesis [62]. Thus N-glycan pathway can potentially contribute to the resistance to apoptosis, a hallmark property of cancer (Fig.4B).

### Other O-glycan biosynthesis

POFUT1, RFNG and ST6GAL1 are involved in the biosynthesis of the fucose linked o-glycans that are known to regulate Notch signaling pathway. So as expected among the direct interactors of these proteins we find NOTCH1 and NOTCH2. Notch signaling pathway has strong implications for cancer and it is known regulate cell proliferation and chemoresistance [63]. In addition, ST6GAL1 is known to glycosylate MUC4 [64] and likely also other mucins with a functional interaction has been suggested (STRING). As mentioned earlier mucins have been linked to growth factor signaling and evasion of apoptosis (see above). Thus, this O-glycan biosynthetic pathway can potentially contribute to two hallmark properties of cancer viz. sustained growth and evasion of apoptosis, which may include chemoresistance (Fig.4B).

### GPI anchor biosynthesis

GPI biosynthetic genes were found among both glyco-oncogenes and glyco-TSGs. *PGAP3, PIGQ, PIGS, PIGX* and *PIGZ* were among the glyco-oncogenes while *PIGG, PIGL, PIGN* and *PIGV* were among the glyco-TSGs. GPI anchor biosynthesis genes present among the glycogenes were involved in the biosynthesis of the core saccharide unit of the GPI anchor (*PIGQ, PIGX, PIGZ*) as well as its attachment to the protein (*PIGS*). PGAP3 catalyzes the removal of unsaturated fatty acid from the GPI anchor in the Golgi. While the GPI anchor biosynthetic genes present among glyco-TSGs were mainly involved in the modification of the core saccharide with the addition of phosphate (*PIGG, PIGN*) or deacetylation of the N-acetlyglucosamine residue (*PIGL*). *PIGV* present in glyco-TSGs encodes for a mannosyltransferase involved in the biosynthesis of the core saccharide part of the GPI anchor. To understand how this pathway may contribute to oncogenesis, we looked at their interactors. There was no significant interaction between GPI biosynthetic pathway and the cancer census genes.

### Characterization of *B4GALT5* as a glycooncogene

The gene encoding GOLPH3 is amplified in several solid tumors and has been shown to act as an oncogene [17]. We recently showed GOLPH3 regulates the proteostasis of B4GALT5, by acting as an adaptor to promote the incorporation of B4GALT5 in COPI vesicles at the Golgi apparatus [19]. This action prevents the lysosomal transport of B4GALT5 and thus prevents its degradation. In cancers where GOLPH3 levels are increased, B4GALT5 levels are also increased and GSL production is increased. This increase in B4GALT5 and the consequent change in GSL levels are essential for the oncogenic action of GOLPH3 [19]. So, we hypothesized that *B4GALT5* itself may act as an oncogene if its copy number is amplified.

To experimentally evaluate the oncogenic activity of *B4GALT5*, a Class III glycooncogene, we first measured the rate of proliferation of cells expressing B4GALT5. NIH3T3 Cells expressing *B4GALT5* proliferated faster compared to control cells as measured by CFSE assay **(Fig.5A)** demonstrating that B4GALT5 expression promotes cell proliferation. Besides increased cell proliferation key cellular properties that contribute to carcinogenesis are absence of contact inhibition of growth and the anchorage independent proliferation of cells. We monitored these two properties in NIH3T3 cells expressing B4GALT5 by using focus formation assay and soft agar assay respectively. NIH3T3 cells transfected with *GOLPH3* and *RAS*^*Val12*^ mutant were used as positive controls. NIH3T3 cells expressing *B4GALT5* showed more colonies in anchorage-independent soft agar assay compared to pcDNA3.1 (empty vector) transfected control cells and at levels comparable to that observed following GOLPH3 or RAS^*Val12*^ overexpression **(Fig.5B)**. In a similar fashion, cells expressing B4GALT5 also showed increased number of foci compared to GFP transfected control cells and again at levels comparable to that observed with cells overexpressing GOLPH3 or RAS^*Val12*^ mutant **(Fig.5C)**. Thus, increased expression of B4GALT5 confers oncogenic properties including increased rate of proliferation, anchorage independent growth and reduced contact inhibition of growth.

**Figure 5:**
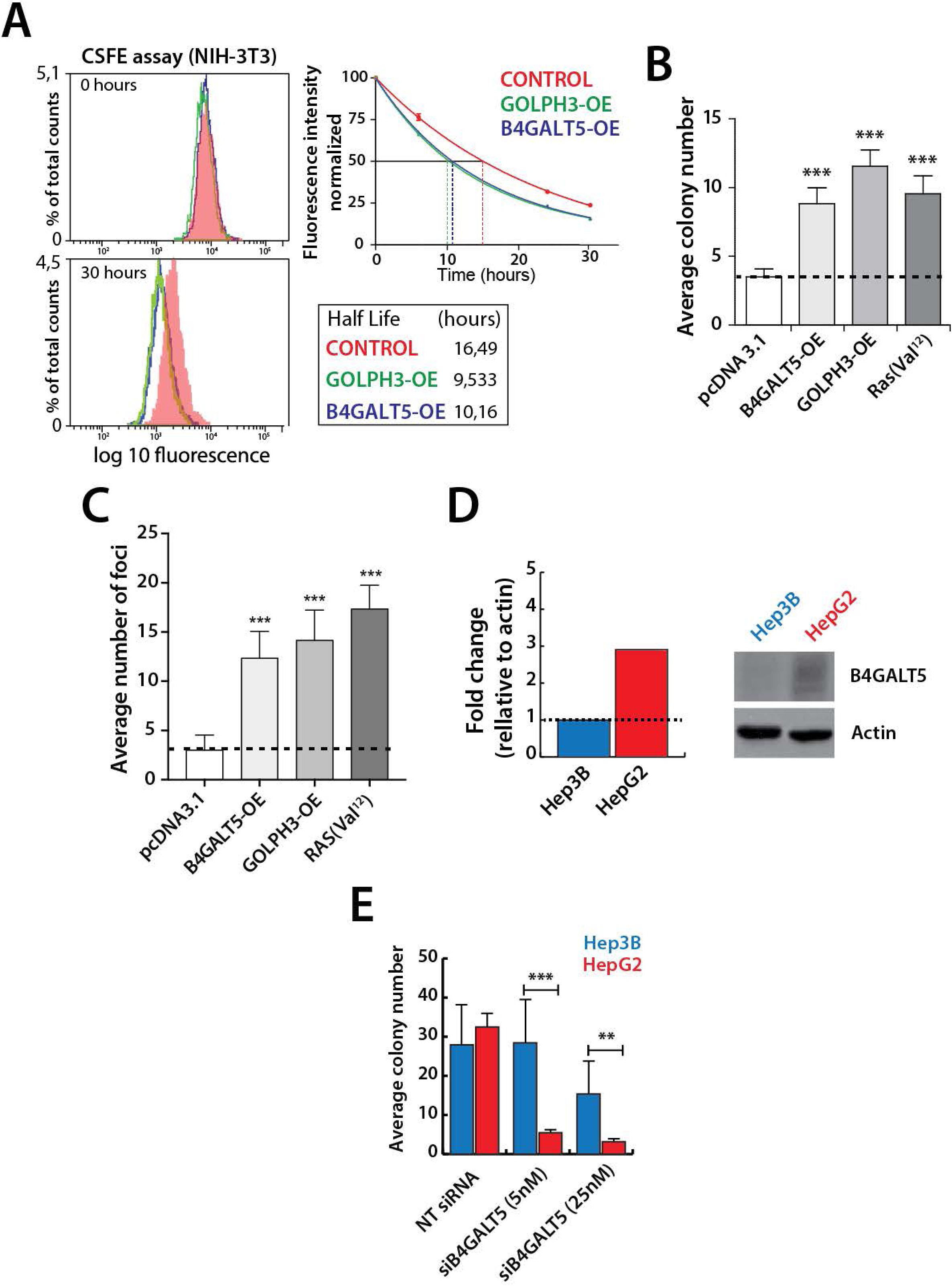
B4GALT5 overexpression promotes oncogenic growth and proliferation *in vitro*. **(A)** *In vitro* proliferation assay in NIH3T3 cells. B4GALT5 overexpressing cells were incubated with CFSE dye at different time points as indicated. Cells were washed and processed for flow cytometry analysis using BD FACS ARIA III cell sorter. The histogram (left) and graph (right) representing normalised mean fluorescence intensity (MSI) over time in cells overexpressing B4GALT5 and GOLPH3 used as a positive control. Data plotted from at least three biological replicates. **(B)** Soft agar colony assay depicting oncogenic role of B4GALT5 in anchorage-independent growth. Graph representing average number of colonies obtained from NIH3T3 cells overexpressing either B4GALT5 or GOLPH3 and RAS^Val12^ as a positive control. Data plotted are means ± SEM of at least three independent experiments; p-value significance calculated by student T-test; *** p-value <0,001. **(C)** Foci formation assay in NIH3T3 cells overexpressing either B4GALT5 or GOLPH3 and RAS^Val12^ depicting contact inhibition growth. Quantification graph showing the average number of foci obtained from different treatments. Data plotted are means ± SEM of at least three independent experiments; *** p-value <0,001. **(D)** Relative mRNA expression of *B4GALT5* across amplified (red) and diploid (blue) liver and ovarian cancer cell lines respectively. Graph represents *B4GALT5* mRNA fold change relative to housekeeping gene *HPRT1* (left). Immunoblot representing relative protein expression of B4GALT5 across amplified (red) and diploid (blue) cancer cell lines of three different origin relative to actin. **(E)** Clonogenic assay in Hep3B and HepG2 cancer cells. Quantification of clonogenic assay in Hep3B and HepG2 cells treated with 5 and 25nM *siB4GALT5*. Data plotted are means ± SD of at least three independent experiments; ** p-value < 0.01 and *** p-value < 0.001.

### *B4GALT5* genetic ablation sensitizes cancer cell growth *in vitro*

The cytoband 20q13.13 containing *B4GALT5* gene is reported to be frequently amplified in several cancers [65], [66], [67]. We also find that *B4GALT5* locus shows significant amplification and a corresponding increase in mRNA levels in 6 cancer types – BRCA, COAD, ESCA, LIHC, STAD and UCS. The frequency of amplification ranged from 2.7% in case of LIHC to 18.3% in case of UCS. Of these cancer types, in LIHC, *B4GALT5* gets classified as a Class III glycooncogene since knock down of *B4GALT5* expression in hepatocellular carcinoma cell lines with amplification of *B4GALT5* locus reduces their growth. Further, in related studies, increased GSL production has been shown to be essential for hepatocellular carcinoma development [68]. So, we have studied the contribution of B4GALT5 to oncogenesis in hepatocellular carcinoma models.

We first identified patient derived hepatocellular carcinoma cell lines with amplification of *B4GALT5* locus based on the genomic data available in the public databases (DepMap project). Two hepatocellular carcinoma cell lines, one with *B4GALT5* amplification (HepG2) and another diploid for *B4GALT5* (Hep3B) were selected. HepG2 showed more mRNA copies and protein expression of B4GALT5 compared to Hep3B **(Fig.5D)**. Together, these data suggested that *B4GALT5* was highly expressed in HepG2 with CNA of *B4GALT5* locus compared to Hep3B with diploid status of *B4GALT5*. Next, to evaluate if the increased expression of B4GATLT5 in HepG2 cells contributes to its oncogenic properties, we tested the effects of *B4GALT5* silencing on cell growth *in vitro*. B4GALT5 expression was silenced in HepG2 and Hep3B cells using siRNAs (along with non-targeting siRNA as control) and their survival monitored by colony forming assays. HepG2 showed an increased sensitivity to reduction in B4GALT5 levels compared to Hep3B cells **(Fig.5E)**. Thus, increased expression of B4GALT5 in HepG2 cells is required for the survival and proliferation of these cells.

### Increased GSL biosynthesis contributes to cell growth in cell lines expressing increased levels of B4GALT5

*B4GALT5* encodes for a key GSL metabolizing enzyme which transfers galactose for a UDP-Gal donor on to GlcCer backbone to form LacCer. Which serves as the precursor for the synthesis of several other complex GSL metabolites. GSLs play a crucial role in cell growth, cell adhesion, cell communication and signalling [69], [41], [70]. So, we measured if increased B4GALT5 expression is associated with an increased GSL production in HepG2 cells. GSLs produced in these cells were monitored using H^3^-Sphingosine pulse chase and HPTLC **[71]**. HepG2 showed high amounts of GSL synthesis compared to Hep3B **(Fig.6A)**. Next, we evaluated if this increased GSL levels is essential for the growth of HepG2 cells. We treated HepG2 and Hep3B cells with known blockers of GSL biosynthesis - Miglustat the growth of these two cell lines were monitored. While Hep3B was nearly insensitive to the treatment with miglustat, HepG2 cells showed a concentration dependent reduction in cell growth (Fig.6B), suggesting that GSL biosynthesis is essential for the growth of HepG2 cells. To test this link between increased GSL production following B4GALT5 overexpression and increased cell growth, we created a catalytically inactive mutant of B4GALT5 by mutating a key tryptophan residue within the UDP-Gal donor pocket conserved among all Gal-T family members (Ref: B4GALT5 ^W296A^). When *B4GALT5*-KO HeLa cells [72] were analyzed by H^3^-Sphingosine pulse chase assay they produced low amounts of GSLs (close of 10% of the Sphingolipids produced in the cell; Fig.6C). We then re-expressed either *B4GALT5*^*WT*^ or the catalytically inactive form *B4GALT5* ^*W296A*^ in these cells and measured the rescue of GSL synthesis. We found *B4GALT5*^*WT*^ showed a rescue of GSL biosynthesis (now 30% of the total sphingolipids produced) compared to control *B4GALT5*-KO cells **(Fig.6C)**. On the other hand, expression of *B4GALT5* ^*W296A*^ did not rescue GSL biosynthesis, where the GSL levels remained the same as in *B4GALT5*-KO cells (**Fig.6C)**. Thus, *B4GALT5*^*W296A*^ mutant is indeed catalytically inactive. Next, we tested whether the catalytic activity of *B4GALT5* was required for expression of oncogenic properties using soft agar assay. *B4GALT5-*

**Figure 6:**
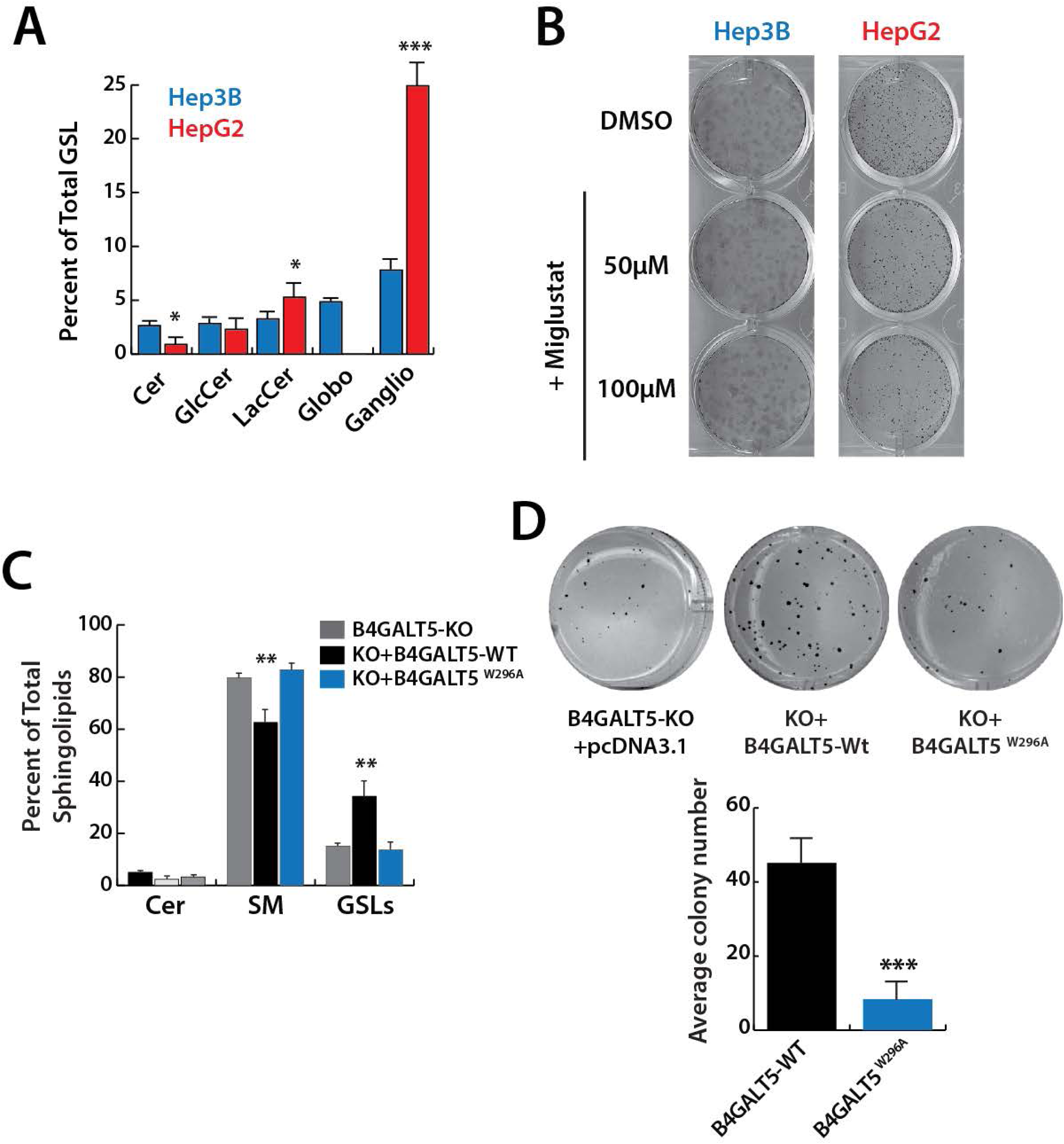
B4GALT5 exerts *in vitro* oncogenic features through GSL biosynthesis. **(A)** Complex glycosphingolipid estimation in Hep3B (blue) and HepG2 (red) cell lines. Graph represents percentage of different complex glycosphingolipids relative to total sphingolipids. Data plotted as mean ± SD of at least three independent experiments; * p-value < 0.05 and *** p-value < 0.001. **(B)** Clonogenic assay in Hep3B and HepG2 cells. Representative image showing the effect on colony formation in Hep3B and HepG2 cells treated with 50 μM and 100 μM of Miglustat (GSL inhibitor). **(C)** Sphingolipid analysis in B4GALT5-KO cells either expressing wild-type B4GALT5 or catalytically inactive B4GALT5^W296A^ mutant; non-transfected B4GALT5-KO cells represented as control. Data plotted as mean ± SD of at least three independent experiments; ** p-value < 0.01 **(D)** Soft agar colony assay in B4GALT5-KO cells expressing either wild-type B4GALT5 or catalytically inactive B4GALT5^W296A^ mutant. Representative images of colonies obtained after 30 days (top). Quantitative graph reporting average number of colonies obtained. Data plotted as mean ± SD from three biological replicates: *** p-value < 0.001.

*KO* cells formed low number of colonies on soft agar plate and expression of B*4GALT5-WT* in these cells increased the number of colonies nearly 10-fold, while expression *B4GALT5*^*W296A*^ did not show a change in the number of colonies compared to B*4GALT5-KO* cells (**Fig.6D)**. Collectively, these findings suggest that increased B4GALT5 levels and the associated increased in GSL levels in cell lines with amplification in B4GALT5 locus contributes to the growth of the cells and to their oncogenic property.

## Discussion

Identification of oncogenes which connect a clear genetic perturbation to a measurable oncogenic process has accelerated the understanding of the oncogenic pathways. While, glycosylation changes have been associated with cancer development and progression for a long time and to the extent that several cancer biomarkers being glycans (for instance CA125, T and Tn antigens) [8] [73], the absence of a strong candidate oncogenes or TSGs among glycogenes has made the exploration of the contribution of glycosylation to cancer an arduous task. So, in order to identify such candidates, we analyzed cancer genome datasets for potential glyco-oncogenes and glyco-TSGs. Our findings here show a potential link between glycan pathways - GSL biosynthesis, proteoglycan biosynthesis, TGN associated lysosomal sorting, GPI anchor biosynthesis and N-glycan biosynthesis and hallmark characteristics of cancers. We identify potential candidate glyco-oncogenes and glyco-TSGs and as a proof of principle elaborate and show the oncogenic properties of B4GALT5, a key enzyme of the GSL biosynthetic pathway. Not only do these findings provide an experimental handle to further explore the mechanistic underpinnings of how glycosylation pathways control oncogenic processes but also open a novel and potential group of enzymes that can be targeted to inhibit carcinogenesis.

In order to identify potential glyco-oncogenes and glyco-TSGs, we have analyzed the genome datasets and mined for potential candidates based on CNA. Several previously studies have analyzed these CNA datasets to identify genes with CNA that are significantly enriched in cancers (see Table S2) or there have been attempts to identify if known oncogenes/TSGs previously identified based on mutations also display CNA (See Table S4 for list of studies). Given that most of the genes in the CNA segment are expected to be passenger genes, the identity of driver genes in these FCNAs remains unclear. Here we developed and implemented a pipeline based on published criteria [31] to analyze TCGA datasets to identify potential driver oncogenes/TSGs. While we have ultimately focused on the glycogenes which is of interest to our laboratories, the prioritized list that is provided with this study (Table S5), we believe, will kick-start the characterization of new potential driver oncogenes/TSGs.

Within top ten oncogenes and TSGs identified in this study nearly half of them are not present in the cancer census gene list, suggesting several interesting candidates’ oncogenes//TSGs may be embedded in our priority list. For instance, CDK12 was classified as a Class II oncogene in CESC. It is not classified as a cancer census gene but there is enough literature pointing towards it role as a likely oncogene [74]. Here we find that amplification of CDK12 in cervical cancer is clinically relevant and thus this study points to classifying CDK12 as an oncogene in CESC. Similarly, PTPN13 a protein tyrosine phosphatase is classified as a Class II TSG in our priority list but is not present in the cancer census gene list. Literature surveys indicate that PTPN13 can act as a potential tumor suppressor gene [75] and our results that it may have strong role as TSG in bladder cancer with clinical relevance points to its importance in bladder cancer.

While several of the pathways identified here have been linked to carcinogenesis earlier including GSL biosynthesis, GAG biosynthesis and N-glycan biosynthesis [76], [77], [78], [79] the study here identifies potential oncogenes /TSGs belonging to these pathways thus consolidating their contribution to cancer development. In addition, it also discovers potentially novel modes of action. For instance, the link of N-glycan pathway to carcinogenesis is mostly associated with the action MGAT5 mediated branching of N-glycans that leads to the production of poly-N-acetyl lactosamine chains that regulate cell surface signaling, specifically the growth factor signaling pathway [80]. On the contrary we here find that downregulation of N-glycan pathway, specifically the early part of the pathway functioning in ER and cis-Golgi involved in biosynthesis of dolichol-anchored glycan biosynthesis and trimming of N-glycans, is associated with carcinogenesis by hosting potential TSGs. Interactome analysis suggested that this pathway may be involved in evasion of apoptosis. Several cancers have also been shown to exhibit impaired N-glycan processing and in ovarian cancer an accumulation high mannose (an indication of inefficient processing in the Golgi) has been documented [81]. Mannosidase downregulation has been shown to increase accumulation of high-mannose glycans in cholangiocarcinoma and this promotes metastatic migration potential of the cancer [82].

Similarly, heparan sulphate biosynthetic pathway has been shown to promote oncogenic signaling due to the role of heparan sulphates in promoting the presentation of the growth factors to their receptors [83]. Here, we find heparan sulphate biosynthetic genes are enriched in glyco-TSGs. Glypicans which are decorated by HS chains have been suggested to act as tumor suppressors at least in some cases [55], [56], possibly depending on the context. Thus, the findings here point to the complexity of action of HSPGs and interpret their actions with caution.

*Contribution of B4GALT5 and GSLs to oncogenesis:* GSLs have long been known as markers of specific cancers. For instance, GD3 is known to present in neuroblastomas while Gb3 marks burkitt’s lymphoma cells. In recent years, it has become clear that GSLs more than being markers of cancers actively contribute to oncogenic signalling. Specifically, the disialylated GSLs like GD3 and GD2 have been shown to promote proliferation of cancer cells while the monosialylated GSLS like GM3 tend to inhibit the cell proliferation. The disialylated GSLs bind to and promote signalling from growth factor receptors and integrins thus promoting growth [41]. We had recently shown that the oncogene GOLPH3 activates Akt signalling and subsequently cell proliferation by promoting increased GSL biosynthesis. A key factor to increase GSL biosynthesis downstream of GOLPH3 is the increased levels of B4GALT5, which seems a rate limiting enzyme in GSL production [19]. Here we find that B4GALT5 itself may act as an oncogene especially in hepatocellular carcinoma. B4GALT5 has been shown to promote growth of gliomas [84, 85] and has also been proposed as a therapeutic target in colorectal cancer [86]. The mechanism of action of B5GALT5 on glioma cancer growth is unclear with a report indicating that the interaction of B4GALT5 with HSC70 may to promote glioma growth [87]. Thus, whether the catalytic activity of the enzyme and hence GSL production contribute to oncogenic signaling is not clear. Here, using drugs targeting GSL biosynthesis and mutant enzymes we show that the oncogenic property of B5GALT5 depends on its catalytic function and hence on the GSLs produced. Further, we also show that in case of hepatocellular carcinoma can act as an oncogene dependent on copy number amplification. Further studies in vivo will shed light on how GSLs altered by increased expression of B4GALT5 contribute to increased growth signaling in hepatocellular carcinoma.

*Limitations of the present study:* For the analysis here, we have concentrated on the CNA and have not considered mutations and epigenetic alterations. While the latter are indeed a potential source for identifying the oncogenes/TSGs we decided to concentrate first on the CNAs to set up a pipeline that can be utilized for genetic alterations. More importantly, CNA of oncogenes have been shown to be better predictors of patient prognosis than the mutated genes [29] and thus starting with CNAs is a pathologically relevant choice. While we did not look for any potentially oncogenic mutations, several mutations that altered the splice site and truncated the protein are present in the database. Whether this contributed to statistically significant presence of mutations that may point to a potential TSG was not analyzed. Further, epigenetic alterations in cancer are well-known to regulate the expression of several critically important genes required for cancer growth and survival [88], [89]. So, exploration the epigenetic landscape of glycogenes in cancer may point to other important oncogenic glycosylation pathways.

## Table legends

**Table 1:** List of cancer types included in the study with their corresponding abbreviations and the number of patient samples analyzed. The TCGA data corresponding to these cancer types were downloaded from cBioportal (https://www.cbioportal.org/).

## References

1 Weinberg R, Hanahan D. The hallmarks of cancer. Cell 2000; 100: 57–70.

2 Hanahan D, Weinberg RA. Hallmarks of cancer: the next generation. cell 2011; 144: 646–674.

3 Varki A. Biological roles of glycans. Glycobiology 2017; 27: 3–49.

4 Pinho SS, Reis CA. Glycosylation in cancer: mechanisms and clinical implications. Nature Reviews Cancer 2015; 15: 540–555.

5 Stowell SR, Ju T, Cummings RD. Protein glycosylation in cancer. Annual Review of Pathology: Mechanisms of Disease 2015; 10: 473–510.

6 Hercules DM. Chemiluminescence resulting from electrochemically generated species. Science 1964; 145: 808–809.

7 Santhanam K, Bard AJ. Chemiluminescence of electrogenerated 9,10-Diphenylanthracene anion radical1. Journal of the American Chemical Society 1965; 87: 139–140.

8 Reis CA, Osorio H, Silva L, Gomes C, David L. Alterations in glycosylation as biomarkers for cancer detection. Journal of clinical pathology 2010; 63: 322–329.

9 Rodrigues E, Macauley MS. Hypersialylation in cancer: modulation of inflammation and therapeutic opportunities. Cancers 2018; 10: 207.

10 Munkley J, Scott E. Targeting aberrant sialylation to treat cancer. Medicines 2019; 6: 102.

11 Pearce OM, Läubli H. Sialic acids in cancer biology and immunity. Glycobiology 2016; 26: 111–128.

12 Ros M, Nguyen AT, Chia J, Le Tran S, Le Guezennec X, McDowall R et al. ER-resident oxidoreductases are glycosylated and trafficked to the cell surface to promote matrix degradation by tumour cells. Nature Cell Biology 2020; 22: 1371–1381.

13 Nguyen AT, Chia J, Ros M, Hui KM, Saltel F, Bard F. Organelle specific O-glycosylation drives MMP14 activation, tumor growth, and metastasis. Cancer cell 2017; 32: 639–653.e636.

14 Guri Y, Colombi M, Dazert E, Hindupur SK, Roszik J, Moes S et al. mTORC2 promotes tumorigenesis via lipid synthesis. Cancer cell 2017; 32: 807–823.e812.

15 Pothukuchi P, Agliarulo I, Russo D, Rizzo R, Russo F, Parashuraman S. Translation of genome to glycome: role of the Golgi apparatus. FEBS letters 2019; 593: 2390–2411.

16 Stanley P, Okajima T. Roles of glycosylation in Notch signaling. Current topics in developmental biology 2010; 92: 131–164.

17 Scott KL, Kabbarah O, Liang M-C, Ivanova E, Anagnostou V, Wu J et al. GOLPH3 modulates mTOR signalling and rapamycin sensitivity in cancer. Nature 2009; 459: 1085–1090.

18 Eckert ES, Reckmann I, Hellwig A, Röhling S, El-Battari A, Wieland FT et al. Golgi phosphoprotein 3 triggers signal-mediated incorporation of glycosyltransferases into coatomer-coated (COPI) vesicles. Journal of Biological Chemistry 2014; 289: 31319–31329.

19 Rizzo R, Russo D, Kurokawa K, Sahu P, Lombardi B, Supino D et al. Golgi maturation-dependent glycoenzyme recycling controls glycosphingolipid biosynthesis and cell growth via GOLPH3. The EMBO Journal 2021; 40: e107238.

20 Dippold HC, Ng MM, Farber-Katz SE, Lee S-K, Kerr ML, Peterman MC et al. GOLPH3 bridges phosphatidylinositol-4-phosphate and actomyosin to stretch and shape the Golgi to promote budding. Cell 2009; 139: 337–351.

21 Isaji T, Im S, Gu W, Wang Y, Hang Q, Lu J et al. An oncogenic protein Golgi phosphoprotein 3 up-regulates cell migration via sialylation. Journal of Biological Chemistry 2014; 289: 20694–20705.

22 Thomas JD, Zhang Y-J, Wei Y-H, Cho J-H, Morris LE, Wang H-Y et al. Rab1A is an mTORC1 activator and a colorectal oncogene. Cancer cell 2014; 26: 754–769.

23 Halberg N, Sengelaub CA, Navrazhina K, Molina H, Uryu K, Tavazoie SF. PITPNC1 recruits RAB1B to the Golgi network to drive malignant secretion. Cancer cell 2016; 29: 339–353.

24 Tan X, Banerjee P, Pham EA, Rutaganira FU, Basu K, Bota-Rabassedas N et al. PI4KIIIβ is a therapeutic target in chromosome 1q–amplified lung adenocarcinoma. Science translational medicine 2020; 12.

25 Chakravarthi BV, Nepal S, Varambally S. Genomic and epigenomic alterations in cancer. The American journal of pathology 2016; 186: 1724–1735.

26 Li W, Olivier M. Current analysis platforms and methods for detecting copy number variation. Physiological genomics 2013; 45: 1–16.

27 Vogelstein B, Papadopoulos N, Velculescu VE, Zhou S, Diaz LA, Kinzler KW. Cancer genome landscapes. science 2013; 339: 1546–1558.

28 Bailey MH, Tokheim C, Porta-Pardo E, Sengupta S, Bertrand D, Weerasinghe A et al. Comprehensive characterization of cancer driver genes and mutations. Cell 2018; 173: 371–385.e318.

29 Smith JC, Sheltzer JM. Systematic identification of mutations and copy number alterations associated with cancer patient prognosis. Elife 2018; 7: e39217.

30 Krištić J, Zoldoš V, Lauc G. Complex genetics of protein N-glycosylation. Glycoscience: biology and medicine 2014: 1–7.

31 Santarius T, Shipley J, Brewer D, Stratton MR, Cooper CS. A census of amplified and overexpressed human cancer genes. Nature Reviews Cancer 2010; 10: 59–64.

32 Gao J, Aksoy BA, Dogrusoz U, Dresdner G, Gross B, Sumer SO et al. Integrative analysis of complex cancer genomics and clinical profiles using the cBioPortal. Science signaling 2013; 6: pl1–pl1.

33 Mermel CH, Schumacher SE, Hill B, Meyerson ML, Beroukhim R, Getz G. GISTIC2. 0 facilitates sensitive and confident localization of the targets of focal somatic copy-number alteration in human cancers. Genome biology 2011; 12: 1–14.

34 Beroukhim R, Mermel CH, Porter D, Wei G, Raychaudhuri S, Donovan J et al. The landscape of somatic copy-number alteration across human cancers. Nature 2010; 463: 899–905.

35 Iorio F, Knijnenburg TA, Vis DJ, Bignell GR, Menden MP, Schubert M et al. A landscape of pharmacogenomic interactions in cancer. Cell 2016; 166: 740–754.

36 Wallace DC. Mitochondria and cancer. Nature Reviews Cancer 2012; 12: 685–698.

37 Bouhaddou M, Eckhardt M, Naing ZZC, Kim M, Ideker T, Krogan NJ. Mapping the protein– protein and genetic interactions of cancer to guide precision medicine. Current opinion in genetics & development 2019; 54: 110–117.

38 Breslow DK, Collins SR, Bodenmiller B, Aebersold R, Simons K, Shevchenko A et al. Orm family proteins mediate sphingolipid homeostasis. Nature 2010; 463: 1048–1053.

39 Bamford S, Dawson E, Forbes S, Clements J, Pettett R, Dogan A et al. The COSMIC (Catalogue of Somatic Mutations in Cancer) database and website. British journal of cancer 2004; 91: 355–358.

40 Kufe DW. Mucins in cancer: function, prognosis and therapy. Nature Reviews Cancer 2009; 9: 874–885.

41 Furukawa K, Ohmi Y, Ohkawa Y, Bhuiyan RH, Zhang P, Tajima O et al. New era of research on cancer-associated glycosphingolipids. Cancer science 2019; 110: 1544.

42 Shiloh Y, Ziv Y. The ATM protein kinase: regulating the cellular response to genotoxic stress, and more. Nature reviews Molecular cell biology 2013; 14: 197–210.

43 Yamaji T, Sekizuka T, Tachida Y, Sakuma C, Morimoto K, Kuroda M et al. A CRISPR screen identifies LAPTM4A and TM9SF proteins as glycolipid-regulating factors. Iscience 2019; 11: 409–424.

44 Meng Y, Wang L, Chen D, Chang Y, Zhang M, Xu J et al. LAPTM4B: an oncogene in various solid tumors and its functions. Oncogene 2016; 35: 6359–6365.

45 Tokuda K, Eguchi-Ishimae M, Yagi C, Kawabe M, Moritani K, Niiya T et al. CLTC-ALK fusion as a primary event in congenital blastic plasmacytoid dendritic cell neoplasm. Genes, Chromosomes and Cancer 2014; 53: 78–89.

46 Argani P, Lui MY, Couturier J, Bouvier R, Fournet J-C, Ladanyi M. A novel CLTC-TFE3 gene fusion in pediatric renal adenocarcinoma with t (X; 17)(p11. 2; q23). Oncogene 2003; 22: 5374–5378.

47 Royle SJ, Bright NA, Lagnado L. Clathrin is required for the function of the mitotic spindle. Nature 2005; 434: 1152–1157.

48 Enari M, Ohmori K, Kitabayashi I, Taya Y. Requirement of clathrin heavy chain for p53-mediated transcription. Genes & development 2006; 20: 1087–1099.

49 Tung K-H, Lin C-W, Kuo C-C, Li L-T, Kuo Y-H, Lin C-W et al. CHC promotes tumor growth and angiogenesis through regulation of HIF-1α and VEGF signaling. Cancer letters 2013; 331: 58–67.

50 Schmid EM, McMahon HT. Integrating molecular and network biology to decode endocytosis. Nature 2007; 448: 883–888.

51 Chi S, Cao H, Chen J, McNiven MA. Eps15 mediates vesicle trafficking from the trans-Golgi network via an interaction with the clathrin adaptor AP-1. Molecular biology of the cell 2008; 19: 3564–3575.

52 Parachoniak CA, Luo Y, Abella JV, Keen JH, Park M. GGA3 functions as a switch to promote Met receptor recycling, essential for sustained ERK and cell migration. Developmental cell 2011; 20: 751–763.

53 Uemura T, Kametaka S, Waguri S. GGA2 interacts with EGFR cytoplasmic domain to stabilize the receptor expression and promote cell growth. Scientific reports 2018; 8: 1–14.

54 Mattera R, Arighi CN, Lodge R, Zerial M, Bonifacino JS. Divalent interaction of the GGAs with the Rabaptin-5–Rabex-5 complex. The EMBO journal 2003; 22: 78–88.

55 Valsechi MC, Oliveira ABB, Conceição ALG, Stuqui B, Candido NM, Provazzi PJS et al. GPC3 reduces cell proliferation in renal carcinoma cell lines. BMC cancer 2014; 14: 1–11.

56 Guo L, Wang J, Zhang T, Yang Y. Glypican-5 is a tumor suppressor in non-small cell lung cancer cells. Biochemistry and biophysics reports 2016; 6: 108–112.

57 Czarnowski D. Syndecans in cancer: A review of function, expression, prognostic value, and therapeutic significance. Cancer Treatment and Research Communications 2021; 27: 100312–100312.

58 Liu Y, Zheng D, Liu M, Bai J, Zhou X, Gong B et al. Downregulation of glypican-3 expression increases migration, invasion, and tumorigenicity of human ovarian cancer cells. Tumor Biology 2015; 36: 7997–8006.

59 Wuyts W, Van Hul W. Molecular basis of multiple exostoses: mutations in the EXT1 and EXT2 genes. Human mutation 2000; 15: 220–227.

60 Santoro M, Moccia M, Federico G, Carlomagno F. Ret gene fusions in malignancies of the thyroid and other tissues. Genes 2020; 11: 424.

61 Hodge JC, Bosler D, Rubinstein L, Sadri N, Shetty S. Molecular and pathologic characterization of AML with double Inv (3)(q21q26. 2). Cancer genetics 2019; 230: 28–36.

62 Liu N, Sun Q, Wan L, Wang X, Feng Y, Luo J et al. CUX1, a controversial player in tumor development. Frontiers in oncology 2020; 10: 738.

63 Aster JC, Pear WS, Blacklow SC. The varied roles of notch in cancer. Annual Review of Pathology: Mechanisms of Disease 2017; 12: 245–275.

64 Zhou X, Kinlough CL, Hughey RP, Jin M, Inoue H, Etling E et al. Sialylation of MUC4β N-glycans by ST6GAL1 orchestrates human airway epithelial cell differentiation associated with type-2 inflammation. JCI insight 2019; 4.

65 Hodgson JG, Chin K, Collins C, Gray JW. Genome amplification of chromosome 20 in breast cancer. Breast cancer research and treatment 2003; 78: 337–345.

66 Wang D, Zhu Z-Z, Jiang H, Zhu J, Cong W-M, Wen B-J et al. Multiple genes identified as targets for 20q13. 12–13.33 gain contributing to unfavorable clinical outcomes in patients with hepatocellular carcinoma. Hepatology international 2015; 9: 438–446.

67 Ptashkin RN, Pagan C, Yaeger R, Middha S, Shia J, O’Rourke KP et al. Chromosome 20q amplification defines a subtype of microsatellite stable, left-sided colon cancers with wild-type RAS/RAF and better overall survival. Molecular Cancer Research 2017; 15: 708–713.

68 Jennemann R, Federico G, Mathow D, Rabionet M, Rampoldi F, Popovic ZV et al. Inhibition of hepatocellular carcinoma growth by blockade of glycosphingolipid synthesis. Oncotarget 2017; 8: 109201.

69 Zhuo D, Li X, Guan F. Biological roles of aberrantly expressed glycosphingolipids and related enzymes in human cancer development and progression. Frontiers in physiology 2018; 9: 466.

70 Yu J, Hung JT, Wang SH, Cheng JY, Yu AL. Targeting glycosphingolipids for cancer immunotherapy. FEBS letters 2020; 594: 3602–3618.

71 Russo D, Della Ragione F, Rizzo R, Sugiyama E, Scalabrì F, Hori K et al. Glycosphingolipid metabolic reprogramming drives neural differentiation. The EMBO journal 2018; 37: e97674.

72 Yamaji T, Hanada K. Establishment of HeLa cell mutants deficient in sphingolipid-related genes using TALENs. PloS one 2014; 9: e88124.

73 Munkley J. The role of sialyl-Tn in cancer. International journal of molecular sciences 2016; 17: 275.

74 Lui GY, Grandori C, Kemp CJ. CDK12: an emerging therapeutic target for cancer. Journal of clinical pathology 2018; 71: 957–962.

75 Freiss G, Chalbos D. PTPN13/PTPL1: an important regulator of tumor aggressiveness. Anti-Cancer Agents in Medicinal Chemistry (Formerly Current Medicinal Chemistry-Anti-Cancer Agents) 2011; 11: 78–88.

76 Li Z, Zhang L, Liu D, Wang C. Ceramide glycosylation and related enzymes in cancer signaling and therapy. Biomedicine & Pharmacotherapy 2021; 139: 111565.

77 Wei J, Hu M, Huang K, Lin S, Du H. Roles of Proteoglycans and Glycosaminoglycans in Cancer Development and Progression. International Journal of Molecular Sciences 2020; 21: 5983.

78 Ahrens TD, Bang-Christensen SR, Jørgensen AM, Løppke C, Spliid CB, Sand NT et al. The role of proteoglycans in cancer metastasis and circulating tumor cell analysis. Frontiers in Cell and Developmental Biology 2020; 8.

79 Taniguchi N, Kizuka Y. Glycans and cancer: role of N-glycans in cancer biomarker, progression and metastasis, and therapeutics. Advances in cancer research 2015; 126: 11–51.

80 Dennis JW, Lau KS, Demetriou M, Nabi IR. Adaptive regulation at the cell surface by N-glycosylation. Traffic 2009; 10: 1569–1578.

81 Chen H, Deng Z, Huang C, Wu H, Zhao X, Li Y. Mass spectrometric profiling reveals association of N-glycan patterns with epithelial ovarian cancer progression. Tumor Biology 2017; 39: 1010428317716249.

82 Park DD, Phoomak C, Xu G, Olney LP, Tran KA, Park SS et al. Metastasis of cholangiocarcinoma is promoted by extended high-mannose glycans. Proceedings of the National Academy of Sciences 2020; 117: 7633–7644.

83 Hassan N, Greve B, Espinoza-Sánchez NA, Götte M. Cell-surface heparan sulfate proteoglycans as multifunctional integrators of signaling in cancer. Cellular Signalling 2020: 109822.

84 Jiang J, Chen X, Shen J, Wei Y, Wu T, Yang Y et al. β1, 4-Galactosyltransferase V functions as a positive growth regulator in glioma. Journal of Biological Chemistry 2006; 281: 9482–9489.

85 Wei Y, Zhou F, Ge Y, Chen H, Cui C, Li Q et al. β1, 4-Galactosyltransferase V regulates self-renewal of glioma-initiating cell. Biochemical and biophysical research communications 2010; 396: 602–607.

86 Chatterjee SB, Hou J, Bandaru VVR, Pezhouh MK, Mannan AASR, Sharma R. Lactosylceramide synthase β-1, 4-GalT-V: A novel target for the diagnosis and therapy of human colorectal cancer. Biochemical and biophysical research communications 2019; 508: 380–386.

87 Sun G, Cao Y, Dai X, Li M, Guo J. Hsc70 Interacts with β4GalT5 to Regulate the Growth of Gliomas. Neuromolecular medicine 2019; 21: 33–41.

88 Lund AH, van Lohuizen M. Epigenetics and cancer. Genes & development 2004; 18: 2315–2335.

89 Chen QW, Zhu X, Li Y, Meng Z. Epigenetic regulation and cancer. Oncology reports 2014; 31: 523–532.

