## Supplementary figures and tables for "Prioritization of driver genes in cancer-associated copy number alterations identifies *B4GALT5* as a glycooncogene"

**Methods:**

**Identification of chromosomal segments with significant CNA:**

Data associated with 32 different cancers in 10845 patients in total were downloaded from http://www.cbioportal.org/ [1]. The data included the putative copy-number values determined using GISTIC 2.0 algorithm [2] for 24776 genes for each cancer patient, relative linear copy-number values for each gene (from Affymetrix SNP6), median expression values, mutation data from whole exome sequencing and clinical data. To identify genes with significant CNA, we developed method inspired by the *Adaptive Daisy Model* (ADaM) [3] that makes use of a statistical model called *Daisy Model* defined in [4] and it was successfully used in [5]. We randomized the GISTIC scores of genes within each chromosome in each patient and the frequency of CNA (a gene was considered to exhibit CNA if GISTIC score >4 for amplification and or GISTIC score <0.5 for deletion) of each gene along the chromosome was calculated. A frequency distribution of the CNA using the randomized score was then plotted. The threshold for significant CNA was then set at the frequency level which covered 99% of this randomized CNA frequency distribution. Any gene with the CNA frequency above the threshold was considered a gene with significant CNA in that cancer type. A contiguous stretch of genes all with significant CNA (with at least 50% overlap in patients) was considered to constitute a chromosomal segment with significant CNA using *Step-wise Constant Fit* algorithm (Regression Analysis and Linear Models: Concepts, Applications, and Implementation, ISBN-13: 978-1462521135). Any chromosomal segment with length >100Mb or < 10Kb were not considered further.

**Prioritization of putative CNA-based oncogenes and TSGs:**

To prioritize genes putative oncogenes starting from CNA information, we followed a so-called *weight-of-evidence* based classification proposed earlier [6]. This classification is defined upon the following set of criteria:

- **Criterion 1:** Significant CNAs (see above)
- **Criterion 2:** Significant change in gene expression correlated with the CNA: Expression levels of genes in patients with CNA was compared to those without. A gene with statistically significant expression change (Wilcoxon test with Benjamini & Hochberg correction, $p<0.05$) in the appropriate direction (increase in case of amplification and decrease in case of deletion) was considered to pass this criterion.
- **Criterion 3:** Correlation between CNA and worsened patient survival: We compared the nonparametric Kaplan-Meier survival curves of patients with and without CNA of a gene and those genes whose CNA is associated with worsened survival (Log-rank test

performed by the *surv_pvalue()* function of the *survminer* CRAN-package) passed this criterion. Of note, the patients were also stage matched (when data was available; Stage I and II were grouped together as early and Stage III and IV were grouped together as late) while doing this comparison.

- **Criterion 4**: Sole identified gene in the chromosomal segment with CNA: If only one gene is present in the chromosomal segment with CNA it passed this criterion.
- **Criterion 5**: Alteration or mutations of other genes in the same pathway: For each gene with significant CNA, we analyzed if other genes belonging to the same pathway (Source: Reactome) were also mutated or have CNA in patients with the same cancer type. If there was a statistically significant (Fisher's test) number of genes belonging to the same pathway were also found to be altered (amplified or having a missense mutation, deleted or having a nonsense/frameshift mutations;) then the gene with CNA passed this criterion.
- **Criterion 6:** Inherited mutations in the genes cause a similar cancer: If any known inherited mutations in the gene leads to the same type of cancer then a gene with significant CNA passed this criterion (data source: Online mendelian inheritance in Man database https://www.omim.org/ ).
- **Criterion 7:** CNA and mutations in the gene are mutually exclusive: If a gene whose CNA and mutation were mutually exclusive it passed this criterion.
- **Criterion 8**: Gene expression change leads to oncogenic features in vitro or in vivo: A gene was considered to pass this criterion if it fulfilled any of the following conditions: a) reduction in expression of a gene in cells with amplification of that gene significantly reduced cell growth/survival compared to cells with no amplification of that gene (data source: DepMAP https://depmap.org/portal/; CERES scores were used comparison and statistically significance calculated by Wilcoxon test, Benjamini & Hochberg, correction $p<0.05$); b) overexpression of a gene increased cell growth or survival [7, 8] and c) transposon-based insertional mutagenesis leads to same type of cancer in animals (data source: Candidate cancer gene database, http://hst-ccgd-prd-web.oit.umn.edu/ ).

The genes were assigned 1 point for each criterion they pass and then they were then classified into 3 Classes with increasing priority based on the number of points gained [6]:

- ***Class IV***, genes passing both Criterion 1 and Criterion 2.
- ***Class III***, Class IV genes with less than or equal to 2 points from other criteria.
- ***Class II,*** Class IV genes with more than 2 points from other criteria.

According to published classification on which this study is based [6], ***Class I*** genes are those that are functional drug targets in clinics. Since we do not have this data, no gene was classified in this group.

**Cell lines and culture conditions:**

Murine NIH3T3 cells were generously gifted by Alessia Varone (Institute of Biochemistry and Cell Biology) and were cultured in DEMEM media supplemented with 10% bovine serum, 2mM L-Glutamine and 100U/ml penicillin and streptomycin. Liver cancer cell lines HepG2 and Hep3B were kind gifts from Nunzio Antonio Cacciola (University of Naples Federico II). HepG2 were cultured in DEMEM media containing 10% FBS, 2mM L-Glutamine and 100U/ml penicillin and streptomycin whereas, Hep3B was cultured in MEM supplemented with 10% FBS, 1mM non-essential amino acid (NAA), 2mM L-Glutamine, 100U/ml penicillin and streptomycin. The HeLa-mCAT8 and TALEN B4GALT5-233 Knock-out cell lines (human cervical cancer cells, female origin) were kind gifts from Kentaro Hanada (National Institute of Infectious Diseases, Tokyo, Japan). Hela-M, Hela-mCAT#8 and B4GALT5-233 KO cells were all cultured in DEMEM media containing 10% FBS, 2mM L-Glutamine and 100U/ml penicillin and streptomycin. All cell lines used in the study was tested and confirmed mycoplasma free and grown under controlled temperature and atmosphere.

**Transfection and RNAi:**

NIH3T3 and Hela-M cells were transfected with either MIRUS or Lipofectamine LTX protocol as per the manufacturer's instructions (in 1% FBS, w/o antibiotics containing media). For obtaining gene silencing in cancer cell lines, a pool of small interfering RNAs (siRNAs) was transfected using Oligofectamine reagent as per manufacturer’s instructions. Two different concentrations, 5 and 25nM of siRNA (pool of 4 different siRNAs) was used in the study.

***In vitro*** **growth and proliferation Assays:**

*Anchorage-independent colony formation assay:*

To test *in vitro* oncogenic transformation, NIH3T3 cells were transfected with pcDNA3.1-B4GALT5-HA, pcDNA3.1-GOLPH3 pcDNA3.1-RASVal12 and empty vector as mentioned above. Approximately, 1 X $10^4$ cells in 2X DEMEM media was diluted with 0.6% (to obtain 0.3% final) sterile agarose and plated gently on top of precoated 0.6% agarose. Wells were supplemented with DEMEM media containing 10% FBS, 2mM L-Glutamine, 100U/ml penicillin and streptomycin and sodium pyruvate on top to avoid drying. Plates were incubated in conventional incubators at 37℃ with 5% atmospheric CO2 for 30 days.

*Clonogenic (Adherent-dependent) assay:*

Briefly, Hep3B and HepG2 cells were cultured in 60mm Petri dishes in respective growth media. The target gene was silenced using a range from 5nM to 50nM siRNA concentration using Oligofectamine transfection protocol mentioned above. After 48 hours of siRNA transfection, cells were trypsinized and counted. An equal number of amplified and diploid cell lines were plated in 12 well plates. To access colony-forming capacities of B4GALT5 amplified cancer cell lines after siB4GALT5, 100 cells were plated and grown for 14 days in low serum conditions. After 14 days, colonies were stained using crystal violet (0.1% w/v) staining and counted using ImageJ.

*Foci formation assay:*

Briefly, NIH3T3 cells were transfected with pcDNA3.1-B4GALT5-HA, pcDNA3.1-GOLPH3 pcDNA3.1-RASVal12 and an empty vector as mentioned above. After 24 hours of transfection, cells were trypsinized and counted before re-plating. Transfected cells were plated in 10 cm culture dishes to obtain high confluency within 24 hours. Cells were maintained in low serum (2.5%) conditions and observed for foci formation for 15-20 days. Foci's were washed with cold PBS, fixed in cold methanol and stained using crystal violet (0.1% w/v) staining for 10 minutes. Stained foci were counted using ImageJ.

**Flow cytometry analysis:**

*Carboxyfluorescein succinimidyl ester (CFSE) proliferation assay:*

NIH3T3 cells after respected treatments were harvested in prewarmed PBS and diluted with equal volume of CFSE dye (2X) and incubated at 37℃ for 10 minutes in dark. Signal was quenched using complete media washed once and resuspended in DMEM containing 5% bovine serum and plated on 6-well plates (50,000 cells/well). A separate plate was prepared with the cells not labelled with CFSE dye (blank). Following an overnight incubation, cells were harvested, washed thoroughly and resuspended in cold PBS. Proliferation analysis was performed using BD FACS ARIA III cell sorter (BD Biosciences).

*Immunoblotting:*

For the estimation of endogenous levels of B4GALT5, amplified and diploid cells were washed twice with PBS and lysed in RIPA buffer (150 mM 299 NaCl, 1% Triton X-100, 0.5% deoxycholic acid, 0.1% SDS, 20 mM Tris-HCl, pH 7.4), supplemented with protease and phosphatase cocktail inhibitor (Roche). The lysates were clarified by centrifugation at high rpm for 5 minutes and quantified using a BCA Protein Assay kit (Pierce). Proteins were resolved on a 12% denaturing polyacrylamide gel and immunoblotted with anti-B4GALT5 antibody containing serum (gift from H.Clausen, University of Copenhagen, Denmark).

**$^{3}$H-Sphingosine pulse-chase and HPTLC:**

Briefly, cells were pulse-labelled with 0.1 µCi/ml (~5nM) $^{3}$H-D-erythro-Sphingosine for 2 hours and chased for 24 hours under optimal confluency. Subsequently, cells were harvested and processed for lipid extraction using an extraction buffer (200µL of 200mM KCL, 500µL methanol, 250 µL chloroform and 250 µLKCL). Finally, lipids were spotted on silica gel high performance-TLC plates (Merck, Germany) and resolved using a mixture of chloroform: methanol: water (65: 25: 4 v/v/v) in a closed glass chamber. After separation, radiolabelled lipids (Cer, GlcCer, LacCer, Gb3Cer, GM3Cer and SM) were analysed using a RITA TLC Analyser (Raytest, Germany) and quantified using GINA (Raytest, Germany) software.

**Real-Time qPCR:**

Cells were harvested washed with PBS and RNA was isolated using RNeasy Mini kit (Qiagen) following the manufactures' protocol. Total RNA was reverse transcribed to cDNA using Quantitative Reverse Transcription kit (Qiagen) as per manufactures protocol. qPCR was performed using target specific primers and LightCycler® 480 SYBR Green I Master Mix (Roche) on a LightCycler® 480 II detection system (Roche). Variation in target gene expression was calculated using comparative CT method (ΔΔCt) normalized against housekeeping gene HPRT1.

**Supplementary Figure legends**

**Figure S1: Glycogenes potentially contain CNA based oncogenes and TSGs. A-B.** Distribution of glycogenes across chromosomes (**A**) or across cytogenetic bands (**B**) shows a near linear correlation with the total number of genes present in a chromosome or cytogenetic band (Pearson's correlation coefficient is indicated). **C**. There were several glycogenes present in the published significantly amplified regions in cancers. The data for the obtained from published studies (see Table X for details). **D**. The expression of several glycogenes is either favorably or unfavorably correlated with prognosis in several cancer types. Data obtained from the human protein atlas (https://www.proteinatlas.org/about/download).

**Figure S2: Identification of statistically significant chromosomal segments with CNA.** A. The distribution of the number of amplified or deleted segments across cancer types. B-C. There is a near linear correlation between the number of amplified and deleted segments across cancer types (B) or chromosomes (C), with the only exception being PRAD which had relatively a greater number of deleted segments (Pearson's correlation coefficient is indicated).

**Figure S3: Characterization of the identified chromosomal segments with CNA. A.** The median length of the identified amplified and deleted chromosomal segments is plotted across cancer types. **B.** The percentage fraction of the already published chromosomal segments with CNA that were also identified in our study was calculated and plotted across cancer types. **C.** The fraction of patients in which an identified chromosomal segment with CNA is present was calculated and plotted against fraction of identified CNA segments with that frequency of alteration.

**Figure S4: Chromosomal segments with pan-cancer alteration in copy number.** The number of cancer types in which a particular chromosomal position shows copy number alteration (amplification in orange and deletion in black) was plotted across the chromosomal length.

**Figure S5: Characterization of chromosomal segments with pan-cancer alteration in copy number. A.** The number of pan-cancer chromosomal segments that were amplified in a chromosome is plotted against the number of pan-cancer chromosomal segments that were deleted in a chromosome. **B.** The number of identified chromosomal segments with pan-cancer alteration was plotted against the length of the chromosome indicates a linear correlation (Pearson's correlation coefficient is indicated).

**Supplementary Table legends:**

**Table S1**: List of glycogenes analyzed in this study.

**Table S2:** The number of published copy number altered regions in each cancer type, the number identified in this study and the overlap.

**Table S3:** Chromosomal segments that were amplified or deleted in more 10 cancer types

**Table S4:** List of known oncogenes and TSGs with CNA.

**Table S5:** List of Class II/III oncogenes and TSGs identified in this study

**References:**


1 Gao J, Aksoy BA, Dogrusoz U, Dresdner G, Gross B, Sumer SO *et al*. Integrative analysis of complex cancer genomics and clinical profiles using the cBioPortal. *Science signaling* 2013; 6: pl1-pl1.

2 Mermel CH, Schumacher SE, Hill B, Meyerson ML, Beroukhim R, Getz G. GISTIC2. 0 facilitates sensitive and confident localization of the targets of focal somatic copy-number alteration in human cancers. *Genome biology* 2011; 12: 1-14.

3 Behan FM, Iorio F, Picco G, Gonçalves E, Beaver CM, Migliardi G *et al*. Prioritization of cancer therapeutic targets using CRISPR–Cas9 screens. *Nature* 2019; 568: 511-516.

4 Hart T, Brown KR, Sircoulomb F, Rottapel R, Moffat J. Measuring error rates in genomic perturbation screens: gold standards for human functional genomics. *Molecular systems biology* 2014; 10: 733.

5 Hart T, Chandrashekhar M, Aregger M, Steinhart Z, Brown KR, MacLeod G *et al*. High-resolution CRISPR screens reveal fitness genes and genotype-specific cancer liabilities. *Cell* 2015; 163: 1515-1526.

6 Santarius T, Shipley J, Brewer D, Stratton MR, Cooper CS. A census of amplified and overexpressed human cancer genes. *Nature Reviews Cancer* 2010; 10: 59-64.

7 Škalamera D, Ranall MV, Wilson BM, Leo P, Purdon AS, Hyde C *et al*. A high-throughput platform for lentiviral overexpression screening of the human ORFeome. *PloS one* 2011; 6: e20057.

8 Lee WJ, Škalamera D, Dahmer-Heath M, Shakhbazov K, Ranall MV, Fox C *et al*. Genome-wide overexpression screen identifies genes able to bypass p16-mediated senescence in melanoma. *SLAS DISCOVERY: Advancing Life Sciences R&D* 2017; 22: 298-308.

### Figure S1

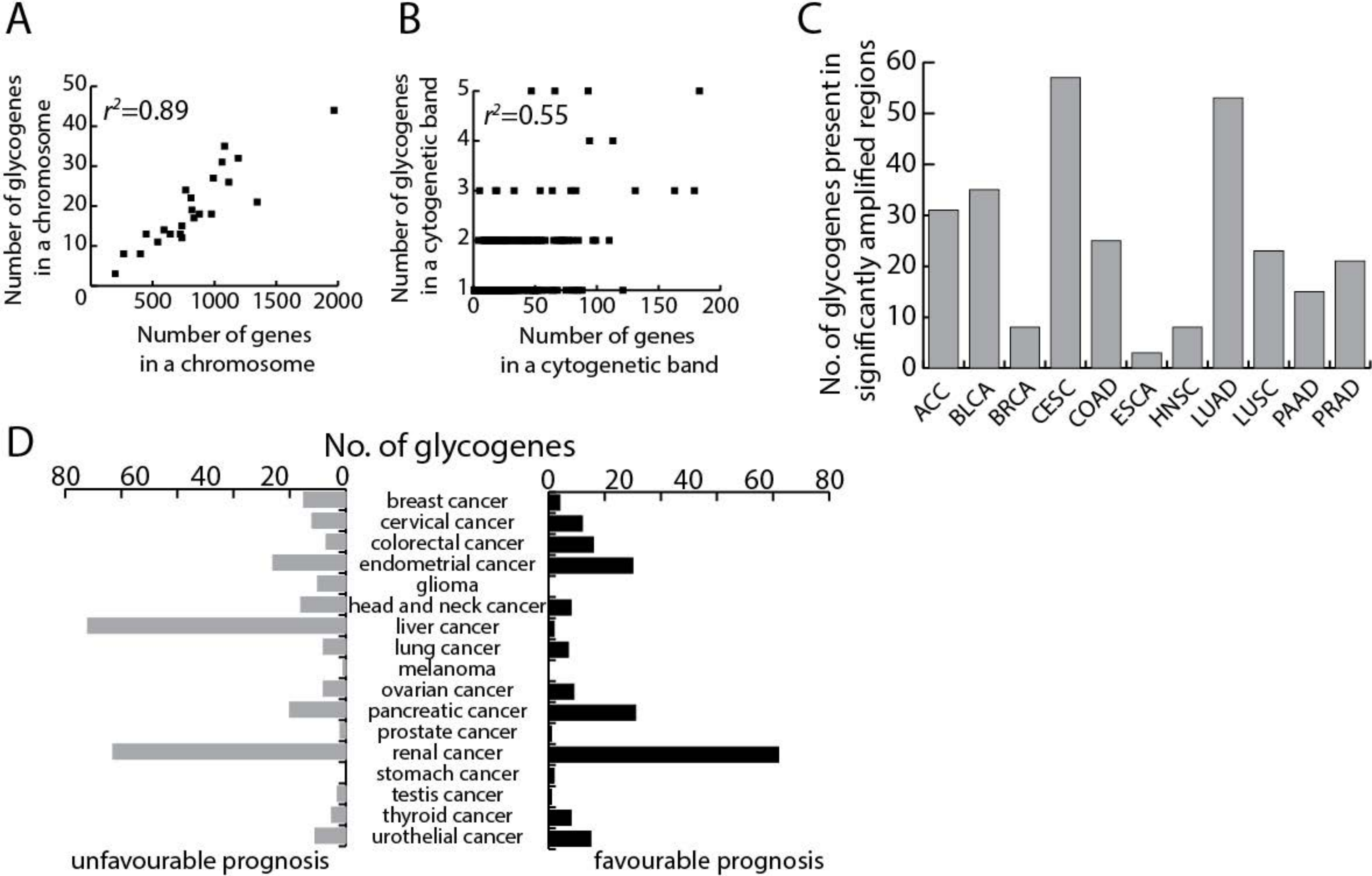
A
r²=0.89
Number of glycogenes in a chromosome
Number of genes in a chromosome
0
10
20
30
40
50
500
1000
1500
2000
B
r²=0.55
Number of glycogenes in a cytogenetic band
Number of genes in a cytogenetic band
1
2
3
4
5
0
50
100
150
200
C
No. of glycogenes present in significantly amplified regions
0
10
20
30
40
50
60
ACC
BLCA
BRCA
CESC
COAD
ESCA
HNSC
LUAD
LUSC
PAAD
PRAD
D
No. of glycogenes
80
60
40
20
0
0
20
40
60
80
breast cancer
cervical cancer
colorectal cancer
endometrial cancer
glioma
head and neck cancer
liver cancer
lung cancer
melanoma
ovarian cancer
pancreatic cancer
prostate cancer
renal cancer
stomach cancer
testis cancer
thyroid cancer
urothelial cancer
unfavourable prognosis
favourable prognosis

Figure S2

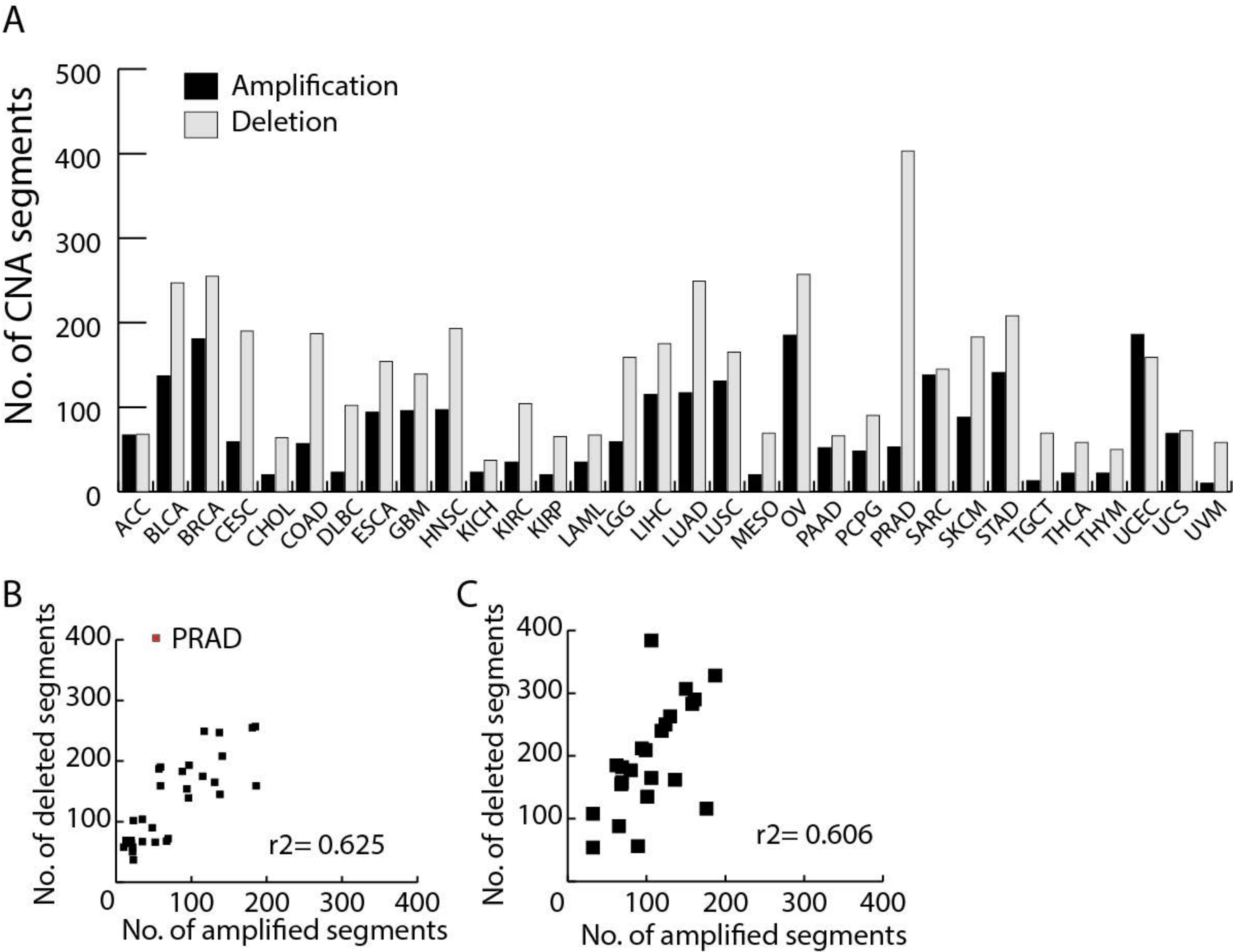
A
B
C
Amplification
Deletion
No. of CNA segments
500
400
300
200
100
0
ACC
BLCA
BRCA
CESC
CHOL
COAD
DLBC
ESCA
GBM
HNSC
KICH
KIRC
KIRP
LAML
LGG
LIHC
LUAD
LUSC
MESO
OV
PAAD
PCPG
PRAD
SARC
SKCM
STAD
TGCT
THCA
THYM
UCEC
UCS
UVM
PRAD
No. of deleted segments
No. of amplified segments
r2= 0.625
r2= 0.606
0
100
200
300
400

### Figure S3

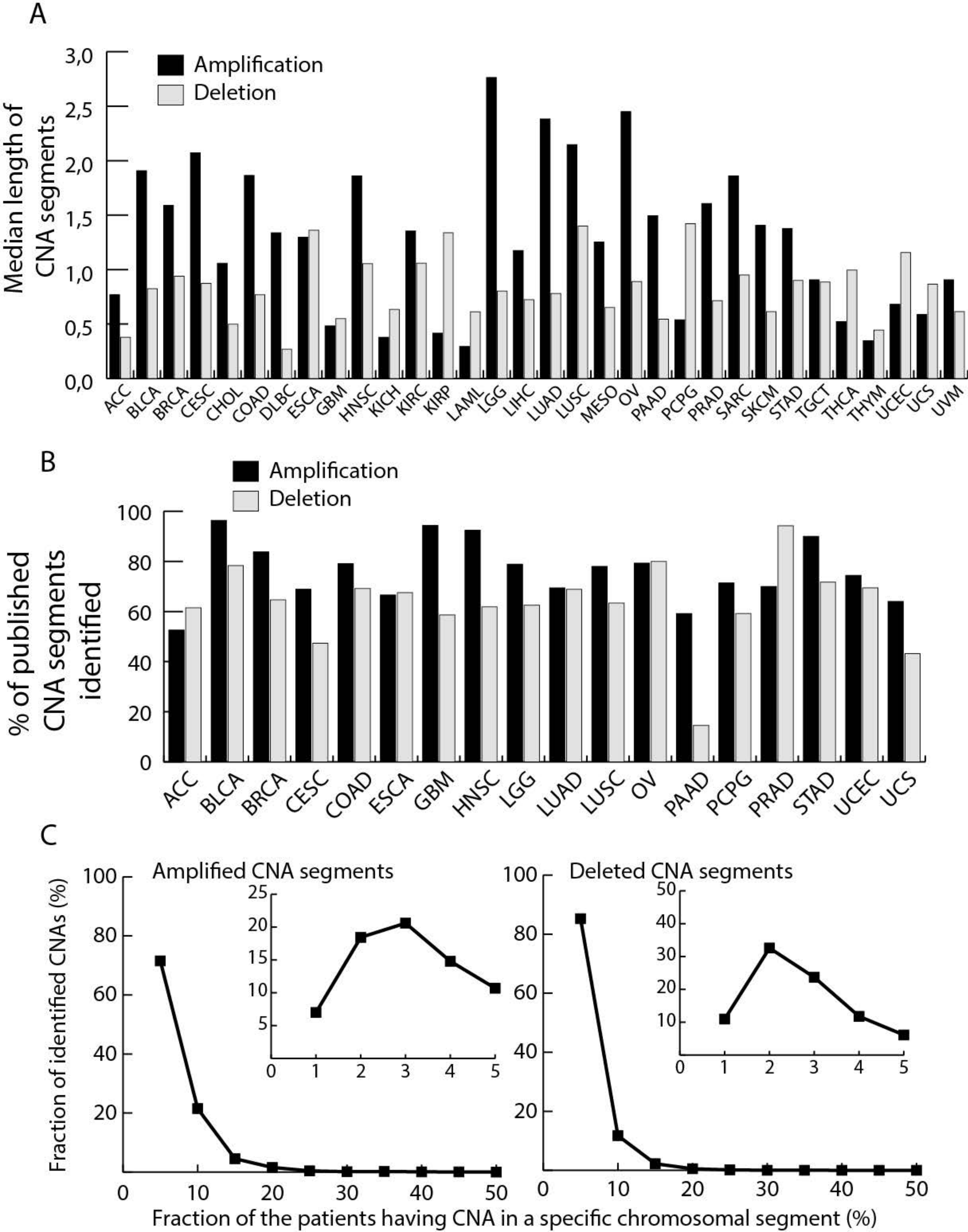
A
Amplification
Deletion
Median length of CNA segments
3,0
2,5
2,0
1,5
1,0
0,5
0,0
ACC BLCA BRCA CESC CHOL COAD DLBC ESCA GBM HNSC KICH KIRC KIRP LAML LGG LIHC LUAD LUSC MESO OV PAAD PCPG PRAD SARC SKCM STAD TGCT THCA THYM UCEC UCS UVM
B
Amplification
Deletion
% of published CNA segments identified
100
80
60
40
20
0
ACC BLCA BRCA CESC COAD ESCA GBM HNSC LGG LUAD LUSC OV PAAD PCPG PRAD STAD UCEC UCS
C
Amplified CNA segments
Deleted CNA segments
Fraction of identified CNAs (%)
100
80
60
40
20
0
10
20
30
40
50
25
15
5
Fraction of the patients having CNA in a specific chromosomal segment (%)

Figure S4

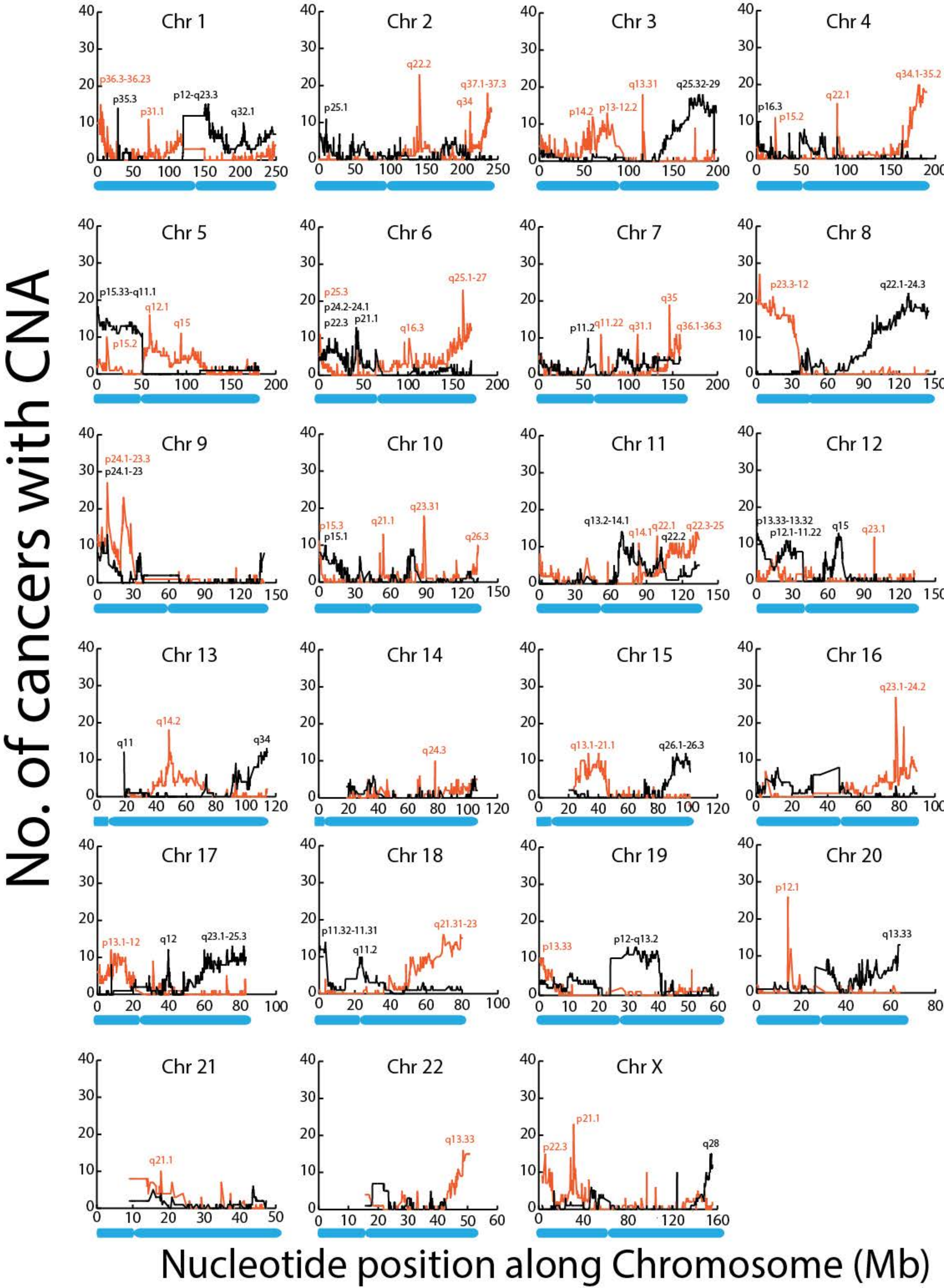
No. of cancers with CNA
Nucleotide position along Chromosome (Mb)
Chr 1
p36.3-36.23
p35.3
p31.1
p12-q23.3
q32.1
Chr 2
p25.1
q22.2
q37.1-37.3
q34
Chr 3
p14.2
p13-12.2
q13.31
q25.32-29
Chr 4
p16.3
p15.2
q22.1
q34.1-35.2
Chr 5
p15.33-q11.1
p15.2
q12.1
q15
Chr 6
p25.3
p24.2-24.1
p22.3
p21.1
q16.3
q25.1-27
Chr 7
p11.2
q11.22
q31.1
q35
q36.1-36.3
Chr 8
p23.3-12
q22.1-24.3
Chr 9
p24.1-23.3
p24.1-23
Chr 10
p15.3
p15.1
q21.1
q23.31
q26.3
Chr 11
q13.2-14.1
q14.1
q22.1
q22.2
q22.3-25
Chr 12
p13.33-13.32
p12.1-11.22
q15
q23.1
Chr 13
q11
q14.2
q34
Chr 14
q24.3
Chr 15
q13.1-21.1
q26.1-26.3
Chr 16
q23.1-24.2
Chr 17
p13.1-12
q12
q23.1-25.3
Chr 18
p11.32-11.31
q11.2
q21.31-23
Chr 19
p13.33
p12-q13.2
Chr 20
p12.1
q13.33
Chr 21
q21.1
Chr 22
q13.33
Chr X
p22.3
p21.1
q28

### Figure S5

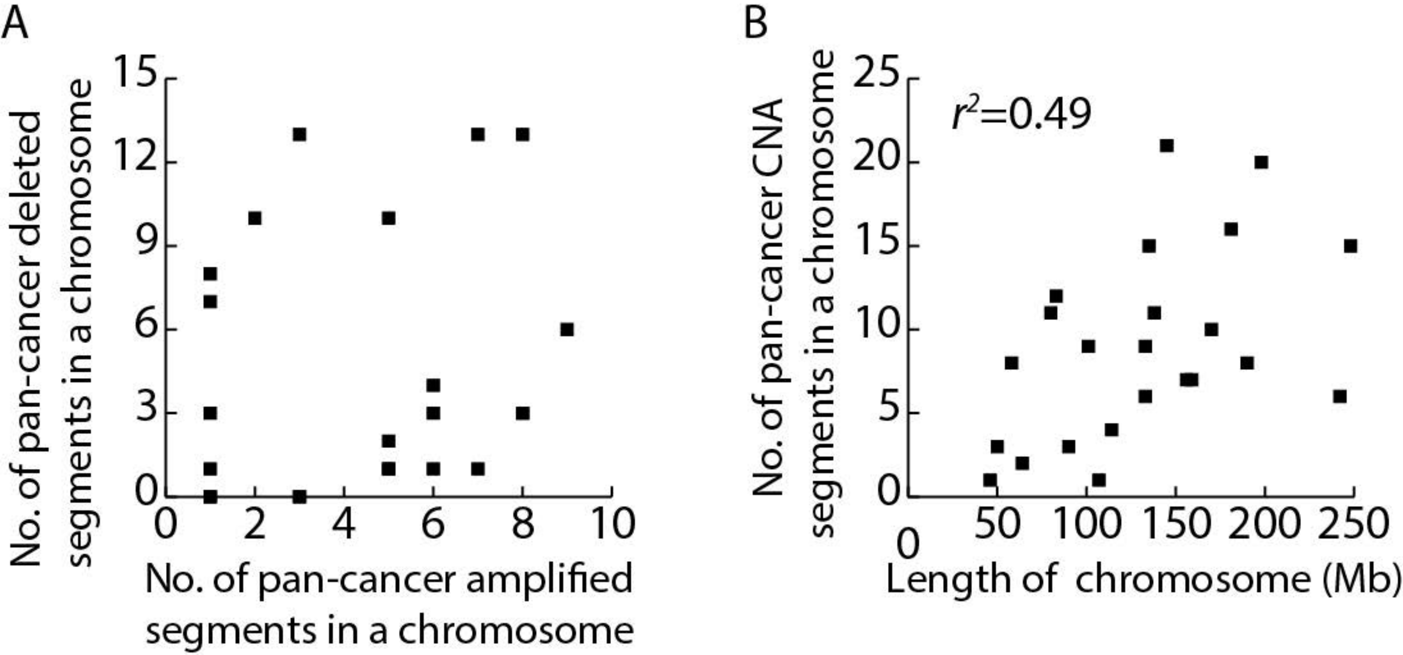
A
B
No. of pan-cancer deleted segments in a chromosome
No. of pan-cancer amplified segments in a chromosome
0
3
6
9
12
15
0
2
4
6
8
10
No. of pan-cancer CNA segments in a chromosome
Length of chromosome (Mb)
r2=0.49
0
5
10
15
20
25
50
100
150
200
250
